# Multi-city analysis of synergies and trade-offs between urban bird diversity and carbon storage to inform decision-making

**DOI:** 10.1101/2024.06.21.600078

**Authors:** R.P. Kinnunen, C.D. Ziter, B. Frei

## Abstract

Cities are particularly vulnerable to the impacts of biodiversity loss and climate change. Urban greenspaces are important ecosystems that can conserve biodiversity and help offset the carbon footprint of urban areas. However, despite large-scale tree planting and restoration initiatives in cities, it is not well known where trees or vegetation should be planted or restored to achieve multiple benefits. We considered urban greenspaces as nature-based solutions for urban climate mitigation and biodiversity conservation planning. Using bivariate mapping, we examined the spatial synergies and trade-offs between bird functional diversity and carbon storage in ten Canadian cities spanning a gradient of geography and population, and modelled the relationships between vegetation attributes and both bird diversity and amount of carbon. We found carbon and biodiversity are weakly positively correlated across the ten cities, however, this relationship varied in strength, direction and significance. Our maps highlight areas within our target cities where greenspaces could be managed, restored, or protected to maximize carbon storage and conserve biodiversity. Nationwide, our results also show that forest management strategies that promote increases in canopy cover and the proportion of needle-leaved species in urban greenspaces are potential win-win strategies for biodiversity and carbon. Our study shows NbS strategies are not always generalizable across regions. National policies should guide municipalities and cities using regional priorities and science advice, since a NbS promoting biodiversity in one region may, in fact, reduce it in another.

## Introduction

Biodiversity loss and the catastrophic effects of climate change on communities worldwide are among the most pressing threats to humans and nature today (IPBES, 2019; Ripple et al., 2020). Although fundamentally interlinked, global strategies to address these challenges have been separate, with growing recognition that integrated approaches are needed (Pettorelli et al., 2021; Turney et al., 2020). Nature-based solutions (NbS) that aim to protect, manage and restore ecosystems while promoting human well-being and biodiversity have been proposed as potential solutions to synergistically address biodiversity and climate crises (Seddon et al., 2020, 2021).

Cities are particularly vulnerable to the impacts of climate change and biodiversity loss (Pörtner et al., 2022). The effects of high temperatures and other extreme weather and climate-related events (e.g., flooding, reduced air quality due to forest fires) are felt more acutely and affect more people in urban compared to rural areas (USGCRP, 2018). Cities thus play a central role in global and national sustainability initiatives, reflected in Sustainable Development Goal 11; Sustainable Cities and Communities (United Nations, 2022) and the Kunming-Montréal Global Biodiversity Framework Target 12; Enhance Green Spaces and Urban Planning for Human Well-Being and Biodiversity (https://www.cbd.int/gbf/targets/12). Planting, restoration, conservation, and management of urban forests are examples of NbS employed by policymakers to combat the impacts of climate change and biodiversity loss in cities while creating economic, environmental, social, and public health benefits for people (Pinto et al., 2022). During the past 20 years, cities worldwide have launched ambitious, often federally funded, large-scale tree-planting and ecosystem restoration initiatives, frequently with climate change mitigation as their driving goal. These include New York City’s Million Trees initiative (Campbell, 2014) and City of Montréal’s Urban Forest Action Plan, aiming to plant 500,000 trees by 2030 (Ville de Montréal, 2020). Other planting and restoration initiatives are underway in cities worldwide (FAO, 2018).

These initiatives can have many positive environmental and health impacts, but may not deliver these without strategic planning (Sousa-Silva et al., 2023), such as ensuring restoration and conservation take place with the greatest environmental and societal impact (Gann et al., 2019; Ordóñez & Duinker, 2013). This is important as poorly planned and executed tree planting or restoration projects can increase carbon emissions and have harmful long-term impacts on biodiversity, ecosystems and human well-being (Di Sacco et al., 2021; Lewis et al., 2019). However, it can be challenging to determine where and how trees should be planted, restored or protected to maximize benefits not only for climate change mitigation but for multiple goals. For example, in temperate forests, maximizing forest carbon stocks by changing the number and arrangement of trees through thinning or planting may benefit some elements of biodiversity but negatively affect others (Sabatini et al., 2019). Poorly planned initiatives can also have unplanned consequences if e.g., native biodiversity is excluded by introduced tree species with the potential to become invasive (Kull et al., 2019). In cities, tree planting initiatives can also be biased toward deciduous species, even where native conifers have been shown to support biodiversity (Fontana et al., 2011). Tree planting and restoration should be viewed as part of a multidisciplinary decision-making process that carefully considers trade-offs and synergies between desired outcomes (Brancalion & Holl, 2020).

Urban greenspaces, including street trees, public parks, golf courses, and residential gardens, represent important ecosystems that can conserve biodiversity and help offset the carbon footprint of urban areas, mitigating some effects of climate change (Churkina, 2016). Greenspaces vary in the ecosystem services or resources they provide, as the structure and composition of vegetation can differ greatly depending on the type of greenspace (Hutt-Taylor & Ziter, 2022; Threlfall et al., 2016). For example, structurally varying vegetation sequesters more carbon (Gough et al., 2019) and provides more ecological niches supporting species-rich and functionally diverse animal communities (Davies & Asner, 2014). Biodiversity-rich, well-functioning urban greenspaces also improve people’s connection to nature, provide aesthetic enjoyment and recreational opportunities, and enhance people’s mental and physical well-being (Fuller et al., 2007; Taylor & Hochuli, 2015).

Birds are great indicators of urban biodiversity and ecosystem function as they are responsive to environmental change, easy to observe, and their occurrence is associated with specific characteristics of the ecosystem (Fraixedas et al., 2020; Kinnunen et al., 2022; Morelli et al., 2021). Birds are also a highly speciose and diverse group, and the distinctive characteristics of each bird species provide a direct link to ecosystem functioning (Cadotte et al., 2011): birds act as ecosystem engineers, seed dispersers, disease regulators, pollinators, nutrient depositors, and invertebrate and rodent predators (Morante-Filho & Faria, 2017). Bird diversity in urban greenspaces is generally explained by greenspace characteristics including area, vegetation structural complexity, or canopy cover (Chamberlain et al., 2007; Fernández-Juricic, 2000; Kang et al., 2015; La Sorte et al., 2020). However, differences in vegetation composition may also be important. The balance of coniferous and deciduous woody plants, the presence of more native conifers, and the variation in canopy cover often positively influence birds (Dyson, 2020; Fontana et al., 2011; Schütz & Schulze, 2015).

While many studies examine bird diversity in urban greenspaces, multi-city studies are still relatively rare (but see Callaghan et al., 2019; Jokimäki et al., 2018). Further, studies often focus on larger cities (Chin et al., 2022; Melles et al., 2003; Smith et al., 2014), yet these results may not always translate to smaller cities (Norton et al., 2016). Synergies and trade-offs between bird diversity and ecosystem functioning are also understudied (but see Belaire et al., 2022; Di Marco et al., 2016; Girardello et al., 2019) despite the importance of such an approach (Seddon et al., 2020). Locating carbon- and biodiversity-rich greenspaces in cities would allow NbS to be implemented in these areas to ensure carbon remains stored and biodiversity is maintained, enabling targeted conservation measures. As the overall aim of many current funding programs and conservation and restoration efforts is to provide multiple benefits (e.g., conserving biodiversity and climate change mitigation), a clearer understanding of the synergies and trade-offs between multiple measures of bird diversity, carbon storage and vegetation attributes in urban greenspaces would enable better urban conservation and restoration decisions.

Here we consider urban greenspaces as a nature-based solution for climate mitigation and biodiversity conservation planning. Using bivariate mapping, we first examine the spatial synergies and trade-offs between bird functional diversity, an indicator of ecosystem resilience, and forest carbon storage by mapping bird and carbon hotspots in ten Canadian cities. We then ask how greenspace attributes (mean and standard deviation (SD) of canopy cover, SD of tree stand age and height, and proportion of broad-leaved trees) affect patterns of bird diversity (species diversity, species richness, and functional diversity), as well as the amount of carbon stored in greenspaces nationwide and regionally (across cities with similar vegetation attributes). We generally predict that bird diversity and carbon storage will increase as canopy cover increases (Chen et al., 2024; Villaseñor et al., 2021) and that other vegetation attributes will have a varying effect on bird diversity and carbon (see Table 1 for predictions). Our results can help urban land managers select sites for conservation, restoration, or management promoting greener cities for multiple goals. As ecosystems that have more functionally diverse bird communities are generally more resilient and better able to withstand the impacts of climate change (Marjakangas et al., 2022), knowledge of how bird functional richness varies in different greenspaces is an important component of building equitable and sustainable cities. By identifying areas with high carbon storage potential, cities can better address the impacts of climate change. Finally, our results show the generalizability of NbS strategies across cities to inform national or provincial programs and policies.

**Table 1.** Our predictions for the relationships between urban greenspace vegetation and bird diversity or carbon storage.

| Vegetation variable | Effect on biodiversity | Effect on carbon | References |
| --- | --- | --- | --- |
| Increase in canopy cover | Increase in bird diversity | Increase in carbon | (Chen et al., 2024; Villaseñor et al., 2021) |
| High canopy cover SD<sup>1</sup> | Lower bird diversity, as higher fragmentation may enable only edge specialists to inhabit the area | Lower carbon, as gaps in canopy cover in urban habitats likely correspond to housing, roads, grass or other similar land use | (Fernández-Juricic, 2001) |
| High tree stand age SD | Higher bird diversity, as these areas likely support more functionally diverse bird communities | Lower carbon, as taller and older trees tend to store more carbon | (Bae et al., 2018; Fernández-Juricic, 2000; Gray et al., 2016) |
| High tree stand height SD | Higher bird diversity, as these areas likely support more functionally diverse bird communities | Lower carbon, as taller and older trees tend to store more carbon | (Bae et al., 2018; Fernández-Juricic, 2000; Gray et al., 2016) |
| Increase in the proportion of broad-leaved trees | Decrease in bird diversity, as urban greenspaces and forests with more conifers generally have a positive impact on bird communities | Increase in carbon storage, as deciduous species tend to be the most dominant street trees in the United States and Canada and thus likely to store the most carbon. | (Clapp et al., 2014; Dyson, 2020; Tilghman, 1987) |

<sup>1</sup> SD = Standard deviation

## Methods

We compiled our dataset using publicly available community science and satellite data. We collected data for ten Canadian cities: Victoria, BC, Vancouver, BC, Calgary, AB, Regina, SK, Winnipeg, MB, Thunder Bay, ON, Windsor, ON, Toronto, ON, Montréal, QC, and Halifax, NS, distributed geographically across the country (Fig 1). We chose cities that vary in population size and land area (Table 2).

**Figure 1.**
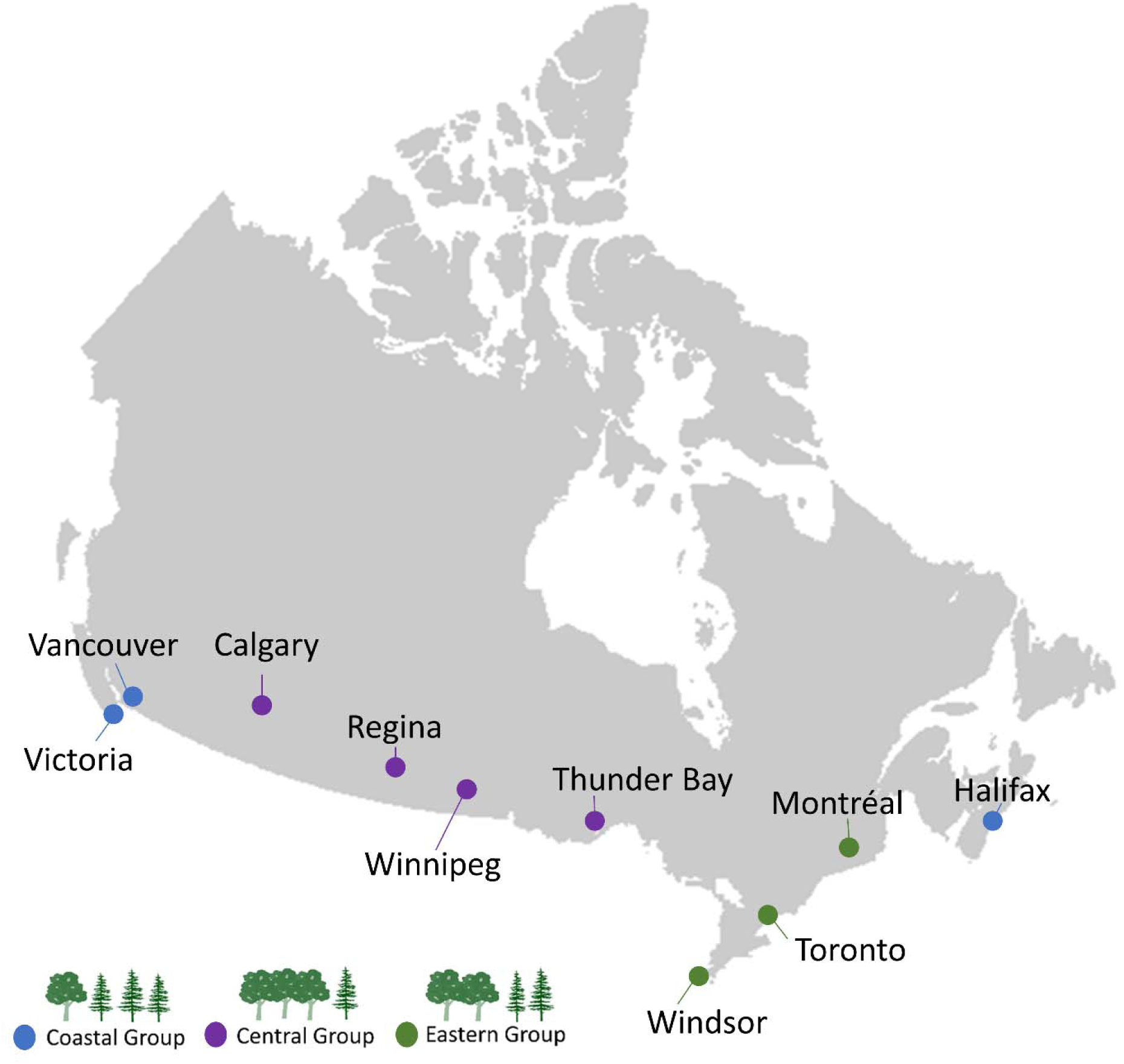
Map showing the Canadian cities used to investigate the relationships between forest attributes and bird diversity and carbon storage, and the spatial similarity of bird functional richness and carbon storage. The different colors show the grouping of cities with similar vegetation based on PCA results.

**Table 2.** The provinces, population sizes and land areas (km^2^) (from Statistics Canada’s 2021 Census) of the ten Canadian cities included in the analyses, as well as the number of bird and carbon sites (grids) per city.

| City | Province | Population size | Area (km<sup>2</sup>) | # Bird Sites | # Carbon Sites |
| --- | --- | --- | --- | --- | --- |
| Victoria | British Columbia | 397,237 | 695.29 | 103 | 710 |
| Vancouver | British Columbia | 2,642,825 | 2,878.93 | 164 | 2179 |
| Calgary | Alberta | 1,481,806 | 5,098.68 | 91 | 327 |
| Regina | Saskatchewan | 249,217 | 4,323.66 | 10 | 139 |
| Winnipeg | Manitoba | 834,678 | 5,285.46 | 95 | 394 |
| Thunder Bay | Ontario | 123,258 | 2,550.79 | 21 | 394 |
| Windsor | Ontario | 422,630 | 1,803.17 | 29 | 233 |
| Toronto | Ontario | 6,202,225 | 5,902.75 | 417 | 2877 |
| Montréal | Quebec | 4,291,732 | 4,670.10 | 306 | 3296 |
| Halifax | Nova Scotia | 465,703 | 7,276.22 | 67 | 1713 |

### eBird data

To identify bird diversity hotspots, we downloaded the eBird Basic Dataset for Canada from eBird.org (Sullivan et al., 2014). eBird is an online bird observation database that provides high-quality bird abundance and distribution data collected by community science volunteer observers. The volunteers submit bird sighting checklists to a central repository, listing the species seen on one bird-watching occasion. Checklists are either complete checklists, listing all birds detected and identified, or incomplete checklists, omitting some observed species. Regional filters identify outliers based on sighting coordinates allowing editors to assess uncommon species sightings, ensuring high data quality (Wood et al., 2011). Our compiled data consisted of species observations and species abundance (number of individuals for each species). We filtered the eBird dataset using bash scripts and R Statistical Software (version 4.2.2; R Foundation for Statistical Computing, Vienna, Austria). We selected only checklists that fell within a census metropolitan area bounding box defined by Statistics Canada and only complete checklists to account for unreported species sightings (Sullivan et al., 2014). Additionally, we selected only checklists that followed specific protocols (stationary, random, traveling, traveling-property specific, area), had 10 or fewer observers, covered a distance of 1 km or less, and had a survey time between 5 and 240 minutes (Johnston et al., 2021). These filters allowed us to refine data and standardize effort between checklists to account for different survey effort across cities. We selected observations recorded between April 1st and October 31st between 2011 and 2021 to match vegetation data and the leaf-bearing season in northern regions. As our main aim was to identify bird diversity hotspots within urban greenspaces, we discarded observations of waterfowl (Anseriformes), as waterfowl species sightings inland are generally flyovers and not indicative of species using urban greenspaces.

We used R package *KnowBR* (Lobo et al., 2018) to assess the survey completeness of eBird species inventories across each city. *KnowBR* calculates survey coverage per unit space as the final slope of the relationship between the number of species observed and the number of database entries, that are used as a surrogate of survey effort (see Appendix S1 for more details). Low-quality survey areas with a slope higher than 0.3, ratio between the number of records and the observed species higher than 3, and completeness lower than 50% were removed from final bird diversity analyses (Lobo et al., 2018).

### Bird functional diversity

We calculated bird functional diversity (FD), or the variation in species’ functional traits reflecting their contribution to the maintenance of different ecosystem processes and functions (Tilman et al., 1997) or species responses to environments (Lavorel & Garnier, 2002). Species-level trait data was extracted from two publicly available databases, the Amniote Life-History database (Myhrvold et al., 2015) and the AVONET (Tobias et al., 2022). We selected 14 traits to cover four underlying functions (habitat use, foraging, movement, and life history), prioritizing traits with direct ecological importance (Appendix S1.3). We ensured each function was represented by at least four traits. Functional diversity was calculated using R package FD (version 1.0-12.1; Laliberté et al., 2014). We calculated three FD indices; 1) Functional richness (FRic), representing the multidimensional convex hull volume occupied by the community, 2) Functional evenness (FEve) measuring the regularity, or evenness, of species in functional trait space weighted by their abundance, and 3) Functional divergence (FDiv), the distribution of abundance within the volume of functional trait space within each community (Laliberté & Legendre, 2010; Villéger et al., 2008; see Appendix S1.4 for descriptions of the FD indices and their ecological meanings). FEve accounts for the relative abundance of species and is independent of species richness (Laliberté & Legendre, 2010).

### Carbon data

We compiled forest carbon biomass data from a recently published high-resolution map of Canada’s terrestrial carbon stocks (Sothe et al., 2022). The authors mapped forest carbon stock in above-ground live biomass, dead plant matter, and below-ground root biomass at 250 m spatial resolution across Canada and produced a spatial carbon distribution map in kg C m^−2^. Briefly, the mapping process used field measurements, satellite data, climate and topographic variables, and a random forest machine learning algorithm (see Sothe et al. 2022 for details). Above-ground live biomass includes all vegetation above the ground (i.e., trees, stems, branches, barks, seeds, flowers, and foliage of live plants).

### Vegetation attributes

We compiled vegetation attributes for cities from Canada’s most recently available forest attributes raster maps, from 2011 (https://open.canada.ca/data/en/dataset/ec9e2659-1c29-4ddb-87a2-6aced147a990). Briefly, the 250 × 250 m resolution maps were produced using the k-nearest neighbors method applied to MODIS satellite imagery and trained from National Forest Inventory (https://nfi.nfis.org) photo plot data (see Beaudoin et al. 2018 for details).

### Data extraction

We assigned georeferenced eBird species records, and carbon and vegetation data to urban areas using population centre boundary files from Statistics Canada (https://www150.statcan.gc.ca/n1/en/catalogue/92-166-X) with a 2 km buffer zone. A population centre is defined as having a population of at least 1000 people and a population density of at least 400 people per square kilometre based on the current census (https://www150.statcan.gc.ca/n1/pub/92-195-x/2021001/geo/pop/pop-eng.htm). To extract data for cities we created a 750 m x 750m raster grid over the population centre shapefiles. We chose this resolution based on the average eBird survey distances for each city, after testing for variation in sampling completeness (%) at four different grid resolutions (250, 500, 750, and 1000 m) across four Canadian cities (see Appendix S1.5). We used the R packages *sp* (version 1.6-0; R. S. Bivand et al., 2013) and *rgdal* (version 1.6-5; R. Bivand et al., 2018) to perform a spatial join between population centres and eBird records. We calculated species richness (total number of species) and Shannon diversity for each grid cell, using R package *vegan* for diversity calculations (version 2.6-4; Oksanen et al., 2022).

We used R package *exactextractr* (version 0.9.1; Baston, 2020) to summarize carbon and vegetation raster values over city grids using the pixel aggregate method. For each polygon, we extracted total forest carbon (kg C/m²), as well as the mean, mode and standard deviation of select vegetation attributes (see Appendix S1 for more information). After testing for correlations between variables (see Appendix S1.6) we chose the mean and standard deviation (SD) of canopy cover, SD of tree stand age, and SD of tree height as our vegetation metrics to be used as explanatory variables in our analyses. We also calculated the proportion of broad-leaved and needle-leaved species in each grid by summing together the total percent composition of both (total trees = total broad-leaved + total needle-leaved species) and then dividing by the percent composition of each group.

### Statistical analyses

All statistical analyses were performed using R Statistical Software (version 4.2.2; R Foundation for Statistical Computing, Vienna, Austria). We used bivariate mapping to illustrate areas of overlap between carbon storage and bird functional richness. We used R package *biscale* (Prener et al., 2022) to calculate k-means clusters at 20% intervals for functional richness and total carbon for each city and assign each raster grid a value of 1–5 depending on the percentile range it belonged to. This created a 5 x 5 matrix with 25 different grid values that varied in importance for carbon and functional richness (Soto-Navarro et al., 2020). We overlayed the bird functional richness map for each city with the carbon map for each city to produce functional richness by total carbon bivariate maps. We considered raster grids with the highest overlapping (the top 20%) values for both functional richness and total carbon in each city as “hotspots”, and grids with the lowest (bottom 20%) values for both as “cold spots” (O’Brien et al., 2023; Soto-Navarro et al., 2020). Grids with the highest values for only functional richness or carbon are considered trade-off areas. To assess the relationship between functional richness and carbon, we calculated Spearman rank correlations between the two.

To investigate how greenspace attributes affect patterns of bird diversity and the amount of carbon nationwide we fit generalized linear mixed-effects models (GLMMs) to data using the function glmmTMB in R package glmmTMB (version 1.1.8; Brooks et al., 2017). Bird diversity (functional richness, functional divergence, functional evenness, species richness, or species diversity) was our response variable, and the select vegetation attributes (mean and SD of canopy cover, SD of tree stand age, SD of tree stand height, and proportion of broad-leaved trees) our explanatory variables. We fit bird FD variables with an ordered beta error distribution and the logit link function (Geissinger et al., 2022), species richness with a negative binomial error distribution and the log link function, and diversity with a Gaussian distribution and the identity link function. We fit total carbon with a Gamma distribution and the log link function. As total carbon is unrelated to bird sampling effort, we did not filter out poorly sampled areas but used the full carbon dataset for these analyses. We thus have comparatively more data for carbon than for bird diversity (Table 2). To make model effects comparable, we scaled data to standardize the range of response and explanatory variables. We included city as a random effect allowing intercepts to vary to account for longitudinal effects and variation across cities. Random intercepts estimate between-group variation in means, as well as variation within groups in each of our response variables. We assessed bias in model fit by visually inspecting residual versus fitted values and used the position of effect sizes and the breadth of confidence intervals to assess relationships (Nakagawa & Cuthill, 2007).

We also wanted to know how bird diversity and carbon vary regionally across cities with similar vegetation attributes. To group cities according to vegetation, we conducted a principal component analysis (PCA) (Appendix S1.7). We standardized data before ordination to account for differences in the units of measurement and used the covariance matrix to perform PCA using the *princomp* function in the *stats* package. We analyzed our data by fitting generalized linear models to data as described above for GLMM analysis but without the added random effect.

## Results

### Bivariate maps

After removing poorly sampled areas from our dataset we had a total of 1303 polygons in ten Canadian cities (Table 2; Fig 1). To test the relationship between functional richness and carbon, we evaluated Spearman rank correlations between the two variables. Across all ten cities, total carbon was positively correlated with bird functional richness (rho = 0.23, *p* < .001; Fig 2). This relationship varied in strength, direction and significance depending on the group (coastal cities group rho = −0.07, *p* = 0.2; central cities group rho = 0.21, *p* = 0.002; eastern cities group rho = 0.27, *p* < 0.001). The bivariate maps showed different spatial overlaps between bird functional richness and carbon across the ten Canadian cities (see Fig 3, Fig 4, and Appendix S2.1-S2.9). There was only a 1.8 % overlap between “hotspots”, that had the highest overlapping values (top 20%) for both functional richness and carbon (bi-class 5-5; Appendix S2.10) and a 3.4 % overlap between top 20-40 % values (bi-classes 5-4 and 4-5; Appendix S2.10). We saw an 8 % overlap between “cold spots”, the clusters with the lowest 20% values for both functional richness and carbon (bi-class 1-1; Appendix S2.10). Our bivariate maps also had a high percentage of areas where bird functional richness was high, but carbon was low (27 % overlap in bi-classes 4-1 and 5-1; Appendix S2.10), but only a very low percentage of areas with high carbon and low functional richness (0.55 %-bi-classes 1-5 and 2-5; Appendix S2.10).

**Figure 2.**
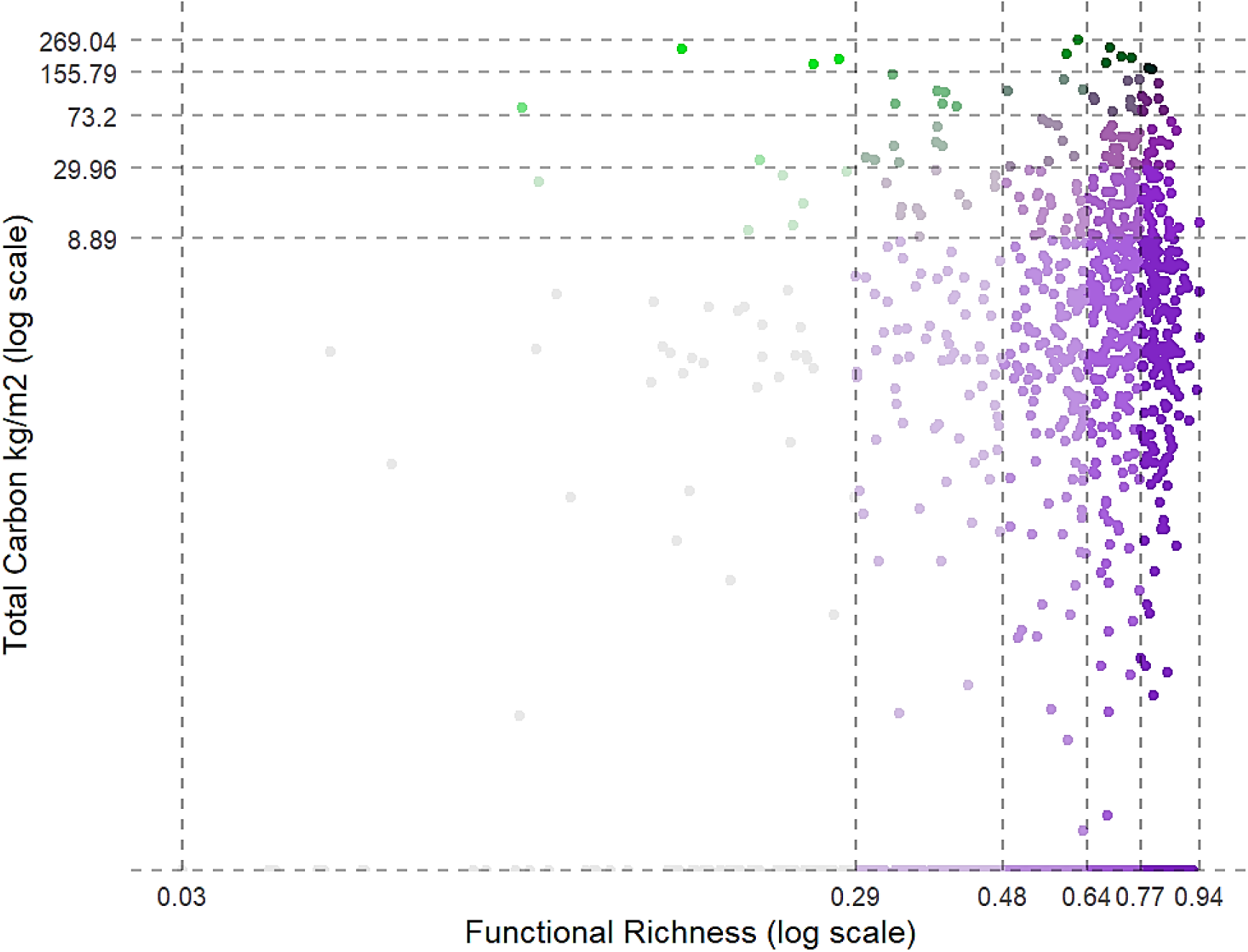
Scatterplot showing the raster grid values for bird functional richness and total carbon for ten Canadian cities. Values are divided into k-means clusters at 20% intervals. The color gradient shows the overlap between different percentile groups for functional richness and carbon. Points in the top right corner (dark green) represent “hotspot” areas with the highest overlapping (the top 20%) values for both functional richness and total carbon. Points in the bottom left corner (light grey) represent cold spots, or grids with the lowest (bottom 20%) values for both variables, and points in the bottom right (dark purple) and top left (bright green) represent “bright spots”, or grids with the highest values for functional richness or carbon.

**Figure 3.**
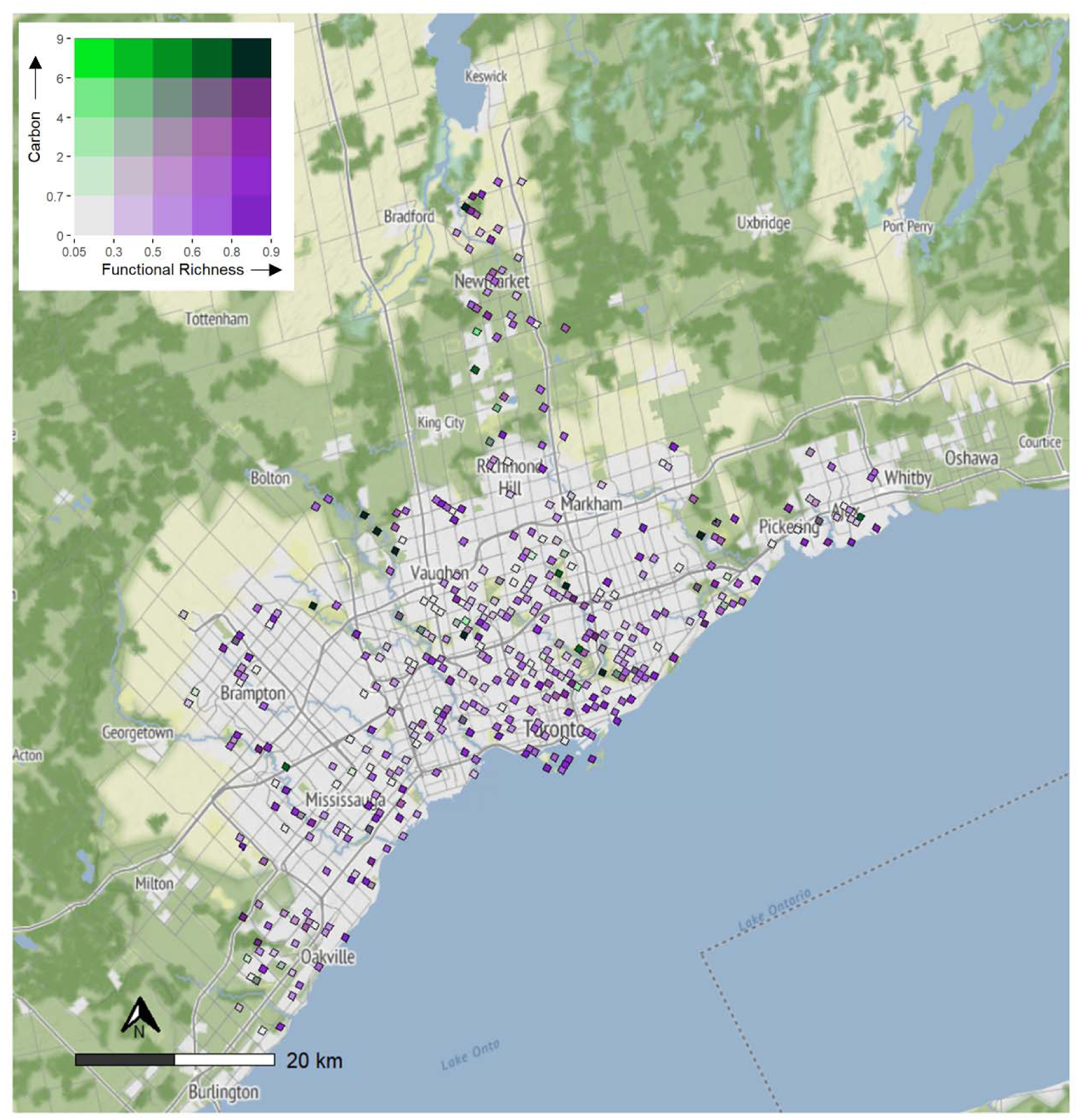
Bivariate map showing the spatial overlap between bird functional richness and above-ground forest carbon (kg C m^−2^) in 750 m grid cells across Toronto, Ontario, Canada. Color scales display k-means clusters at 20% intervals. Dark green regions in the top right corner represent “hotspot” areas with the highest overlapping (the top 20%) values for functional richness and total carbon. Light grey areas in the bottom left corner represent cold spots, with the lowest (bottom 20%) values for both variables. Dark purple regions in the bottom right and bright green regions in the top left represent trade-offs or grids with the highest values for functional richness, or carbon.

**Figure 4.**
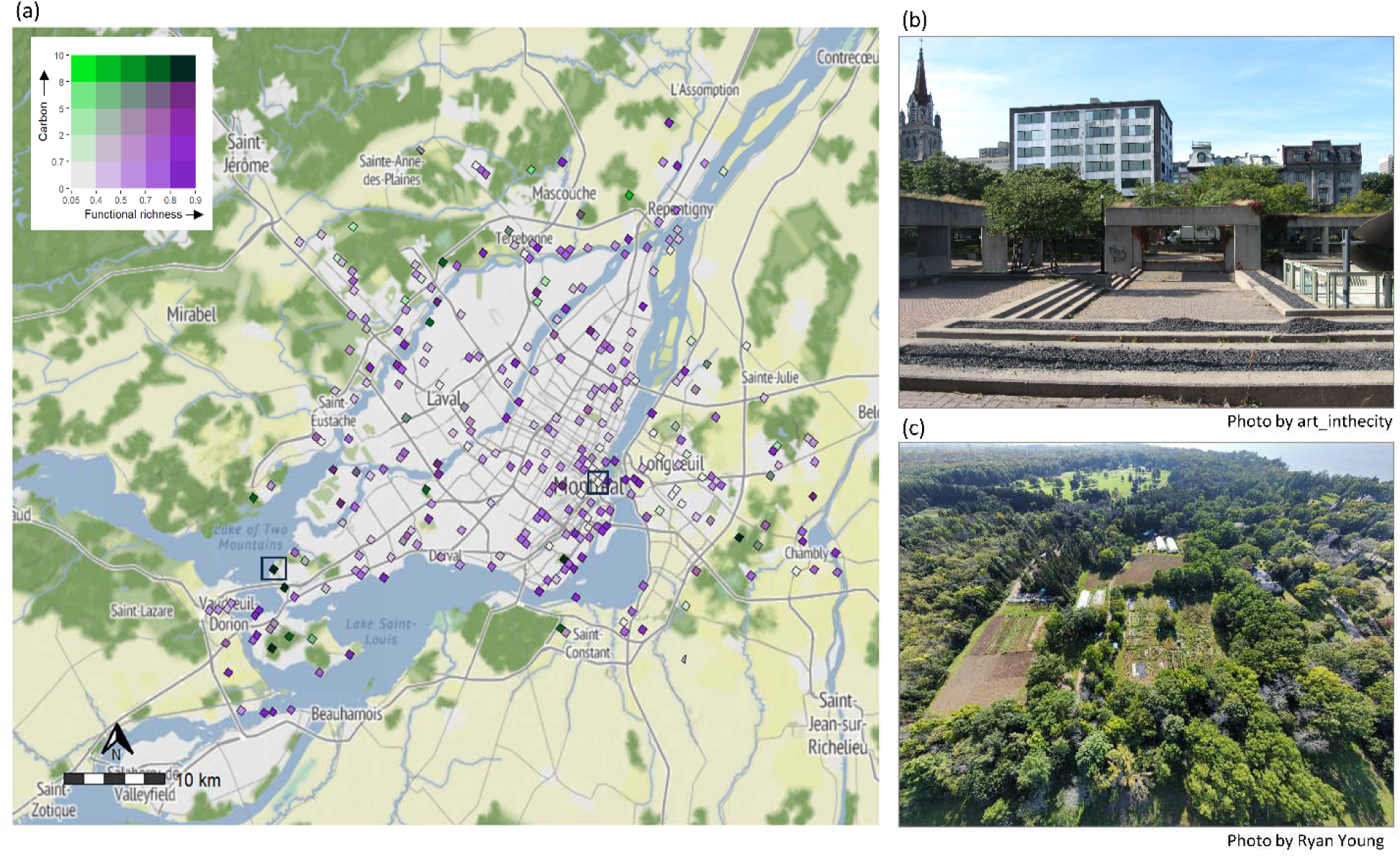
a) Bivariate map showing the spatial overlap between bird functional richness and above-ground forest carbon (kg C m^−2^) in 750 m grid cells across Montréal, Quebec, Canada. Color scales display k-means clusters at 20% intervals. Dark green regions in the top right corner represent “hotspot” areas with the highest overlapping (the top 20%) values for functional richness and total carbon. Light grey areas in the bottom left corner represent cold spots, with the lowest (bottom 20%) values for both variables. Dark purple regions in the bottom right and bright green regions in the top left represent trade-off areas-grids with the highest functional richness, or carbon values. The two grids surrounded by a black box show a cold spot area where carbon and functional richness are both at their lowest (above) and a hotspot area where both values are at their highest (below). Photos visualize the difference between cold spots and hotspots. b) Viger Square is an urban square in Montréal with low functional richness and low carbon (Photo licence CC BY 2.0, https://creativecommons.org/licenses/by/2.0/), and c) Senneville Migratory Bird Sanctuary in the south-west of Montréal, is a 555-hectare protected area containing an array of habitats important for breeding and migrating birds with high functional richness and high carbon.

### Principal component analysis

PC1 explained 62.8% of the total variance, making it a suitable grouping measure. The similarities and dissimilarities in vegetation between cities were visualized in a biplot (see Appendix S1.7). Based on these groupings and the differences in variation in the average broad-leaved species and needle-leaved species percentages across the ten Canadian cities (Appendix S1.7), we clustered cities into three groups (see Fig 1). The first group included the coastal cities of Victoria, Vancouver, and Halifax and had generally high needle-leaved species percentages (18-37%) and low broad-leaved species percentages (5-19%) (Appendix S1.7). The second group included mainly the prairie, or central region cities of Calgary, Regina, Winnipeg, and Thunder Bay with more broad-leaved species (22-44%), low needle-leaved species percentages (6-18%) and generally less canopy cover. The third group included the eastern Canadian cities of Montréal, Toronto, and Windsor. These cities had similar percentages of both broad-leaved (median between 18-32%) and needle-leaved (median between 8-34%) species (Appendix S1.7).

### Data analyses

Across all ten cities, bird functional evenness was positively related to mean canopy cover and negatively related to age SD (Fig 5; see Appendix S2.11. for model estimates). Bird functional richness was positively related to height SD and negatively related to the proportion of broad-leaved species and age SD. Species richness and diversity were positively related to mean canopy cover and height SD and negatively related to the proportion of broad-leaved species and age SD. Total carbon was positively related to mean canopy cover, canopy SD, and age SD and negatively related to the proportion of broad-leaved species and height SD.

**Figure 5.**
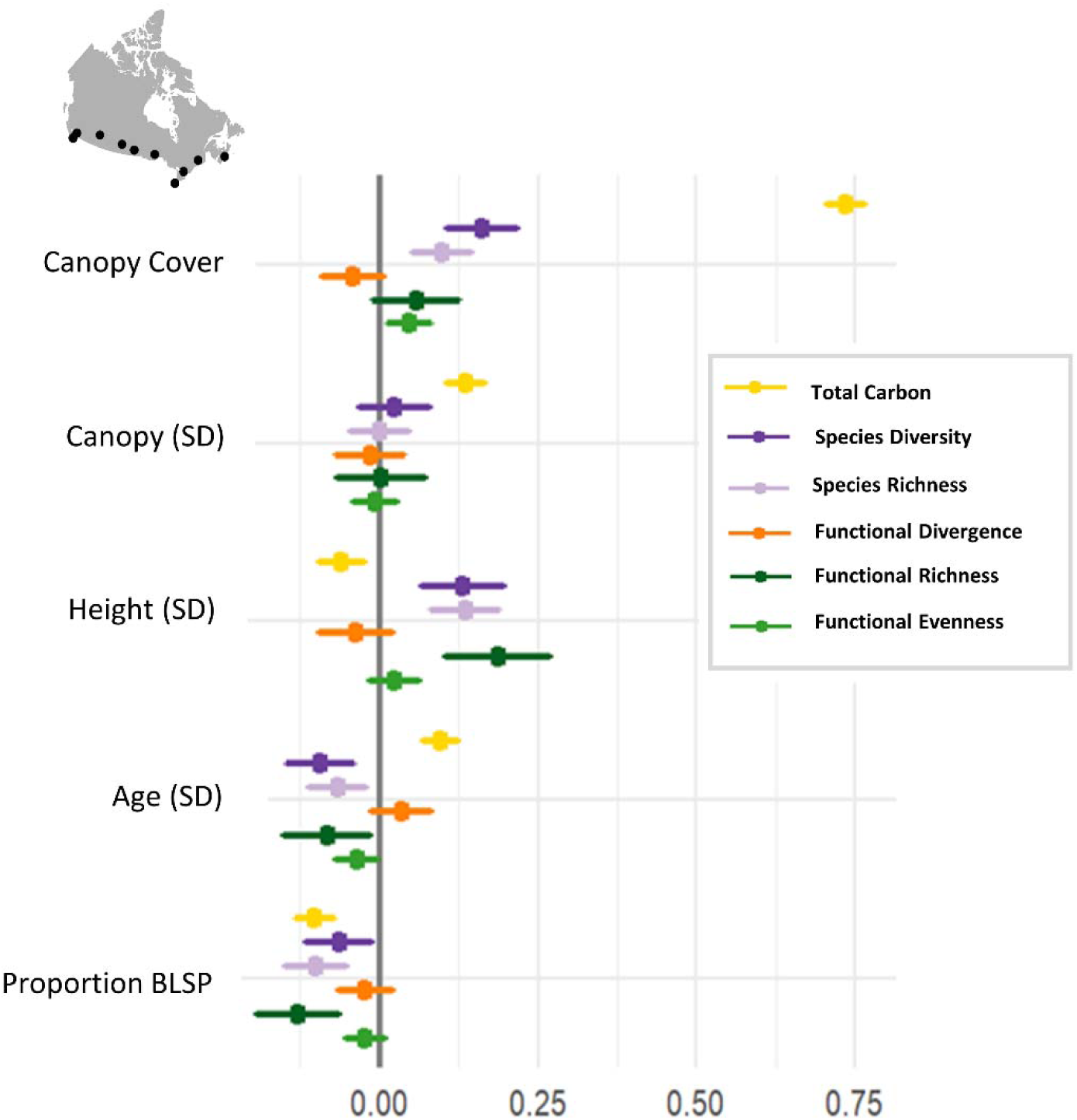
Linear and generalized linear mixed-effects model coefficients showing the relationships between urban greenspace attributes, and bird diversity and carbon variables across ten Canadian cities. Circles represent coefficient estimates, whiskers are 95% confidence intervals. SD = Standard deviation, BLSP = Broad-leaved species.

For the coastal cities group, bird functional evenness and functional richness were positively related to mean canopy cover and height SD and negatively related to age SD (Fig 6; see Appendix S2.12 for model estimates). Bird functional evenness was also negatively related to the proportion of broad-leaved species. Bird functional divergence was negatively related to height SD. Bird species richness and diversity were positively related to mean canopy cover and height SD and negatively related to the proportion of broad-leaved species and age SD. Total carbon was positively related to mean canopy cover and age and height SD and negatively related to the proportion of broad-leaved species. For the central cities group, bird species richness and diversity were positively related to mean canopy cover and the proportion of broad-leaved species (Fig 6). Total carbon was positively related to mean canopy cover and height SD and negatively related to age SD. For the eastern group, bird functional richness was negatively related to the proportion of broad-leaved species (Fig 6). Bird functional divergence was negatively related to height SD and positively related to age SD. Bird species richness and diversity were positively related to mean canopy cover and negatively related to the proportion of broad-leaved species. Total carbon was positively related to mean canopy cover, canopy SD, and height SD.

**Figure 6.**
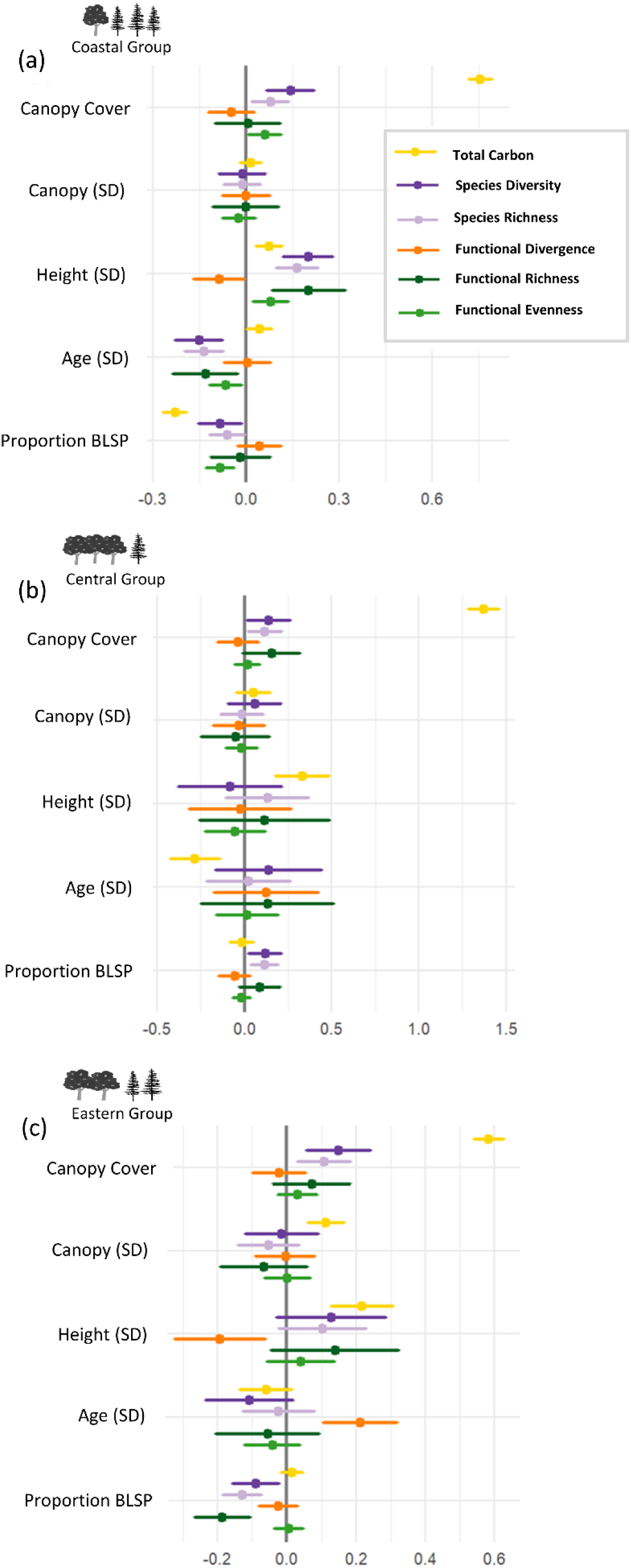
Generalized linear model coefficients depicting the relationships between urban greenspace attributes, and bird diversity and carbon variables for b) coastal group cities c) central group cities and c) eastern group cities. Circles represent coefficient estimates, whiskers are 95% confidence intervals. SD = Standard deviation, BLSP = Broad-leaved species.

## Discussion

In our multi-city investigation, we considered urban greenspaces as a nature-based solution for urban climate mitigation and biodiversity conservation planning. Our bivariate mapping approach identified synergies and trade-offs between bird functional richness and forest carbon storage across Canadian cities that differed in geographic location, size, and human population. Overall, we saw a weak but significant positive correlation between carbon storage and bird functional richness across all ten cities. This relationship differed in strength and direction when cities were grouped by similar vegetation, with the strongest positive correlation seen in the eastern cities of Toronto, Windsor and Montréal, and a weak negative (non-significant) correlation in the coastal group cities of Victoria, Vancouver and Halifax, highlighting the heterogeneous nature of city environments. Hotspots, where both functional richness and carbon were at their highest, were often found in large forest tracts situated in periurban areas or larger city parks (such as Assiniboine forest in Winnipeg, Fish Creek Provincial Park in Calgary, Black Oak Heritage Park in Windsor-part of the Ojibway Prairie park complex, or Parc Angrignon in Montréal; Fig 4, Appendix S2.1-S2.4), or near rivers and ravines (Bois-des-Esprits in Winnipeg, Toronto ravine system; (City of Toronto, 2017); Appendix S2.1). Our hotspots also highlighted the importance of protected areas with diverse habitats specially set aside for conservation, like the Senneville Migratory Bird Sanctuary at the western tip of the Island of Montréal (Fig 4). The sanctuary is part of a new Grand Parc de l’Ouest project in Montréal, that aims to preserve the forests, woodlands, swamps, agricultural lands and ecosystems found within included parks (Ville de Montréal, 2024). Such actions can not only conserve diverse and resilient bird communities but also store high amounts of carbon in line with municipal climate change plans.

Hotspots were overall relatively rare, with “cold spots”, or areas of low carbon and low functional richness being more common within the urban landscape. This is not surprising, considering that cities are often dominated by species tolerant of urbanization that generally have similar characteristics (McKinney, 2006) and that the carbon sequestration and storage potential of many urban greenspaces depends on the intensity of urban land use in the area (Nowak et al., 2013). Cold spots were generally in neighborhoods with small urban squares and parks scattered among residential or industrial areas. One such example is Viger Square in Montréal. It is an urban square with high amounts of impervious surfaces surrounded by high-rise buildings and bordered by major roads on all sides (Fig 4). Notable is that although these areas have low bird diversity and store little carbon, birds still occur in these greenspaces and people observe birds in the area. Cold spots can thus have value in connecting people with nature (White et al., 2023).

Our bivariate maps also showed many areas where bird functional richness was high but carbon was low. These trade-off areas were often found in multifunctional city parks with large sports fields and kids’ playgrounds, surrounded by relatively green suburban areas (e.g., Withrow Park in Toronto, and Daniel Johnson Park in Rivière-des-Prairies–Pointe-aux-Trembles in Montréal; Fig 3, Fig 4). Other functionally diverse areas had very few trees, mainly encompassing large grass fields (Westfort Playfield in Thunder Bay; Appendix S2.5). High bird diversity was thus not only found in areas with extensive urban forests or high canopy cover but also in other habitats. As people’s yards often contain a diverse array of trees and other vegetation including many flowering and fruiting species (Hutt-Taylor & Ziter, 2022), the high bird diversity in these areas may reflect the diverse resources available for birds in suburban areas (Gonçalves et al., 2021; Gray & van Heezik, 2016). It is also possible that the carbon maps used in our study underestimate the amount of carbon storage in residential areas, as field observations using limited number of plots or remote sensing data, both of which were used to create these maps (Sothe et al., 2022), may cause errors in sampling and lead to uncertainties in carbon estimation (Chen et al., 2017; McPherson et al., 2013). As our temporal period considered included both the breeding and migratory seasons, birds may also have been using these neighborhoods and parks as stopover locations. Although migratory birds often prefer heterogeneous forest habitats (Callaghan et al., 2019; Guo et al., 2023) and forest margins with berry-producing shrubs for refuelling during migration (Poirier et al., 2024), our findings suggest that other habitats may also be important for birds.

We also saw many of these high biodiversity-low carbon areas around rivers and coastal areas. These likely highlight the diversity of shorebirds or other waterbirds not excluded from analyses using urban blue spaces to nest or refuel during migration (Appendix S2.13), and may also partly explain the abundance of hotspots near rivers and ravines (discussed above). Although non-passerine species represent a small percentage of total observations (Appendix S2.13), they may influence our functional diversity values and might be driving some of the differences between bird diversity and carbon. However, there is evidence that habitat near urban waterbodies is also important for forest birds and that maintaining these areas can conserve biodiversity (Xie et al., 2022). Additionally, water elements are a common element of urban greenspaces (Konijnendijk, 2024), and considering the functional diversity of all birds visiting and inhabiting these areas is important.

Our results facilitate the development of future studies on drivers of bird diversity and how greenspaces should be managed to maintain existing bird diversity. This could include, for example, not replacing shrubs and other vegetation with trees, even if this would increase carbon storage in the area. If management is done without considering possible trade-offs we risk inadvertently destroying habitat that may benefit birds. For example, planting trees in a grassland may benefit forest species but potentially result in a loss of grassland birds, a group of critical conservation concern (Rosenberg et al., 2019). Future research should also integrate additional taxa of strong conservation interest in cities, such as pollinators and other beneficial insects, which benefit from a variety of habitat types (Ayers & Rehan, 2021; MacInnis et al., 2023).

In contrast, areas with low bird functional richness and high carbon were rare and generally found in larger forest tracts outside of the city (e.g., The Domaine-Seigneurial-de-Mascouche Metropolitan Park near Montréal, Trowbridge Forest near Thunder Bay; Fig 4, Appendix S2.5), but also sometimes adjacent to suburban areas (e.g., Peggy’s Wood in Newmarket, near Toronto; Fig 3). As these forest habitats are undoubtedly important for birds, the results seem paradoxical, particularly as we often see high functional richness in smaller urban greenspaces. The results may reflect the more homogeneous environment in more natural forests or forest plantations with similarly aged trees, likely to attract more specialized birds in contrast to the often more diverse habitat within cities that can increase bird functional richness within urban greenspaces (Hagen et al., 2017). It is also possible that these areas are high human use areas, like in the case of the Domaine-Seigneurial-de-Mascouche Metropolitan Park, part of the Grand-Coteau-Forest-Corridor near Montréal. The park is a popular outdoor location in the region and has many hiking and mountain biking trails and allows dogs on leash (https://mascouche.ca/loisirs-et-parcs/parcs-municipaux/parc-metropolitain). Since human presence alone on forest trails can negatively influence forest bird communities (Bötsch et al., 2018), the high human activity in the area may limit bird diversity.

We were also interested in knowing how greenspace attributes affect patterns of bird diversity and the amount of carbon across the ten cities. Previous studies show that higher canopy cover and structurally complex and heterogenous vegetation are generally positively correlated with bird diversity (Sandström et al. 2006, Ferenc et al. 2014, Threlfall et al. 2016, La Sorte et al. 2020, Callaghan et al. 2021) and our results partly support this. Overall, we show that variation in greenspace vegetation influences the diversity of birds inhabiting these greenspaces. Nationwide, functional evenness was highest in areas with high canopy cover, indicating a more even distribution of traits and implying a more heterogeneous environment supporting diverse bird communities. In contrast, trait distribution was more clumped (lower functional evenness) when there was high variation in tree age within the area, suggesting that species reliant on older trees are found in areas with older trees, and species able to use younger vegetation found in younger neighborhoods with newly planted trees. Bird functional richness was higher in areas that varied in tree height, highlighting the importance of vertical heterogeneity to birds. This is not surprising, as the more layers are available, the more different niches there are for ground, shrub and canopy breeding and foraging birds (Bae et al., 2018; MacArthur & MacArthur, 1961). Our results also showed that bird species richness and diversity were higher in areas with high canopy cover and more variation in tree height, and lower when the proportion of broad-leaved species and variation in tree age was higher. We had predicted that bird diversity would be higher in areas that vary in tree stand age, however, our results did not support our prediction. As greater forest age has been found to increase the abundance of canopy and cavity-nesting birds (Hobson & Bayne, 2000) it is possible that more evenly aged urban forests, particularly if mainly consisting of older trees, may support more birds. Overall, our results imply that higher canopy cover and greater variation in greenspace vertical structure positively influence bird diversity, supporting more bird species and ensuring more resilient and diverse bird communities (Gunderson, 2000). They also suggest that areas with greater variation in tree age and more broad-leaved trees support fewer species overall and that bird communities in these greenspaces are less diverse.

The carbon sequestration and storage potential of urban greenspaces typically depends on greenspace components like canopy cover or plant species present (Wang et al., 2023), the way the greenspaces are managed (Nowak et al., 2013), or the intensity of urban land use in the area (Nowak et al., 2013). We found that the amount of carbon was higher with high and varied canopy cover and greater variation in tree age, and lower in areas that varied greatly in tree height and had more broad-leaved species. Our results support previous studies that found higher carbon storage with higher canopy cover (L. Chen et al., 2024) and show that areas with high canopy cover support both carbon and bird diversity. Our results also show trade-offs between variation in tree stand age and height for birds and carbon. We found that greenspaces with high vertical heterogeneity have high bird diversity but lower carbon storage capacity, while greenspaces with greater variation in tree age store more carbon but have less diverse bird communities. Mature, higher-diameter trees often form the upper canopy and carbon storage is thus generally higher in older tree stands, compared to stands with smaller still-growing trees (Köhl et al., 2017). However, it’s possible that in our case the higher variation in age captures both old high-carbon trees and lower-carbon but higher-density medium-age trees that are more common in cities, leading to higher carbon storage in these areas. In contrast, as greater vertical heterogeneity provides many niches for birds, it may be more important for birds than variation in tree age in urban greenspaces. We also show that as broad-leaved trees increase, both carbon and biodiversity decrease. Our nationwide results suggest that increasing the number of conifers and tree species diversity has the potential to synergistically increase both carbon storage and bird diversity.

Interestingly, these nationwide patterns do not always manifest at a regional or citywide level. When cities are grouped by similar vegetation and climate, the relationships between vegetation and bird diversity and carbon sometimes differ from the nationwide results and vary between groups. Higher canopy cover consistently increased carbon and bird diversity in all three groups. However, there were differences regarding other vegetation attributes. For example, an increase in broad-leaved tree species decreased bird diversity in coastal (Victoria, Vancouver, and Halifax) and eastern group cities (Windsor, Toronto, and Montréal) but increased bird diversity in the central group (Calgary, Regina, Winnipeg, and Thunder Bay). This was likely driven by the lower percentage of conifers in the prairie ecosystem making conifers less ecologically relevant in the region to increase biodiversity (Appendix S1.7). Additionally, tree stand age and height variation influenced carbon storage differently across groups. In the coastal group, carbon storage increased as variation in both age and height increased. In contrast, in the central group variation in tree age decreased, and variation in tree height increased carbon storage, while in the eastern group variation in tree height increased carbon whereas tree stand age had no effect. These differences are likely driven by the differences in vegetation, and vegetation structure, in these cities. The wet coastal climate generally supports more conifers than deciduous trees (Appendix S1.7), like the large, old Western red cedars (*Thuja plicata*) and Douglas firs (*Pseudotsuga menziesii*) found in and around Vancouver. On the other hand, the drier and sunnier central Canadian cities generally have more deciduous trees (Appendix S1.7), like the predominant American elm (*Ulmus americana*) and ash (*Fraxinus americana*) in Winnipeg, while eastern Canadian cities have a mix of deciduous and coniferous trees, such as maples, oaks, hickories, pine trees, and hemlocks.

Our research approach has some limitations and drawbacks. The inclusion of geographically diverse and smaller cities necessitated the use of comparable data products across the country, which is why our vegetation data does not fully temporally match the bird observation or carbon data. We thus assume these have remained comparable. These data products have uncertainties and have been compiled to primarily assess vegetation characteristics or carbon storage in forests outside cities (Beaudoin et al., 2018; Sothe et al., 2022) and are not specifically designed for studies in urban greenspaces. Future research priorities should focus on collecting consistent, national-scale urban data, particularly from urban greenspaces. This would enable more multi-city studies needed to extrapolate results and identify generalizable patterns across cities and continents (Magle et al., 2019). Additionally, our bivariate maps and guidelines do not consider urban social equity. Whiter higher-income neighborhoods often have higher canopy cover than lower-income often multicultural neighborhoods (Myers et al., 2023; Schwarz et al., 2015), and considering where trees are most needed to benefit all communities equitably is an important social consideration when selecting sites for tree planting.

Urban land managers face difficult decisions regarding which sites to conserve, restore, or manage to create greener cities for multiple goals. Our bivariate maps provide a useful guideline for these purposes. As hotspot areas have high biodiversity and high carbon, conserving these areas can help maintain bird diversity and ensure that carbon in these greenspaces remains stored and is not lost over time (O’Brien et al., 2023; Soto-Navarro et al., 2020). Cold spot areas with low biodiversity and low carbon provide opportunities for restoration and tree planting to synergistically conserve biodiversity and mitigate the impacts of climate change. Finally, the trade-off areas identified by our maps can benefit from habitat management and provide further opportunities for research to determine what drives the patterns seen. Our results highlight that forest management strategies that promote increases in canopy cover and the proportion of needle-leaved species in urban greenspaces can be considered win-win strategies for biodiversity and carbon, however, (Gann et al., 2019; Ordóñez & Duinker, 2013) these NbS strategies may not always be generalizable across regions. Tree planting and restoration initiatives should thus be linked at the design stage to strategic urban forest management (Gann et al., 2019; Ordóñez & Duinker, 2013). Our results apply to similar cities worldwide and give an idea of the generalizability of NbS strategies across different geographical ecoregions. These results are also important in a Canadian context, as Canada is a signatory to the Convention on Biological Diversity and the United Nations Framework Convention on Climate Change. As such Canada is committed to NbS to build resilience and help the country meet its 2030 and 2050 climate and biodiversity protection goals and the targets of the Kunming-Montreal Global Biodiversity Framework (ECCC, 2002). Additionally, as populations of many of Canada’s bird species are declining (North American Bird Conservation Initiative Canada, 2019), managing, restoring and planting trees in urban greenspaces in a way that supports bird conservation could directly impact Canada’s biodiversity conservation goals.

Overall, we found synergies and trade-offs between bird diversity, carbon storage and vegetation attributes in urban greenspaces. Our bivariate maps highlight areas where greenspaces could be managed, restored, or protected to maximize carbon storage and conserve biodiversity, enabling better conservation and restoration decisions in our cities. Our results suggest that national policies should be able to guide municipalities and cities using regional priorities, since NbS promoting biodiversity in one region may reduce it in another. This knowledge can inform national, provincial or regional programs and policies.

## Supporting Information

### Appendix S1 Additional methodological details

#### Survey effort

We used R package *KnowBR* (Lobo et al., 2018) to assess the survey completeness of eBird species inventories in the 750 m x 750 m spatial units across each city. *KnowBR* calculates survey coverage per unit space as the final slope of the relationship between the number of species observed and the number of database entries, that are used as a surrogate of survey effort. *KnowBR* calculates the accumulation curve in each spatial unit according to the *exact* estimator (Ugland et al., 2003), as well as performing 200 permutations of the observed data (*random* estimator) to obtain a smoothed accumulation curve which is then fitted to four different asymptotic accumulation functions. These functions can be used to obtain a completeness percentage (the percentage that represents the observed number of species compared to the predicted number of species), which can also be used to estimate the spatial units with likely complete surveys. Low-quality survey areas with a slope higher than 0.3, a ratio between the number of records and the observed species higher than 3, and completeness lower than 50% were removed from final bird biodiversity analyses (Lobo et al. 2018).

#### Functional diversity

We also calculated bird functional diversity (FD). Species-level trait data was extracted from two publicly available databases, the Amniote Life-History database (Myhrvold et al., 2015) and the AVONET (Tobias et al., 2022). The Amniote Life-History database is a systematically compiled database of life-history traits for birds, mammals, and reptiles built for comparative life-history analyses. For species with multiple raw data points, the median value is reported. The AVONET contains comprehensive functional trait data for all birds with both raw data and species averages reported. We accounted for differences in eBird sampling effort by removing poorly surveyed areas, with slope > 0.3 and completeness lower than 50% (Lobo et al. 2018), from our dataset before FD calculations.

We first tested the correlation between all possible continuous traits (n=15) (Appendix S1.1) to avoid trait redundancy (Laughlin, 2014). From strongly correlated (> 0.70) trait pairs we selected only one, leaving us with 8 continuous traits in addition to 6 categorical variables. We then calculated the predictive power score (PPS) between the continuous and categorical trait variables (Appendix S1.2) using R package *ppsr* (version 0.0.2; van der Laken, 2021). PPS is an asymmetric, data type-independent score that can detect linear or non-linear relationships between two columns with a value between 0 (no predictive power) and 1 (perfect predictive power). There were no strong relationships between the continuous and categorical variables (Appendix S1.2), so we selected 14 traits in total to cover four underlying functions (habitat use, foraging, movement, and life history), prioritizing traits that have direct ecological importance (Appendix S1.3). We also ensured that each function was represented by at least four traits. Functional diversity was calculated using R package FD (version 1.0-12.1; Laliberté et al., 2014). We calculated three functional diversity indices; 1) Functional richness (FRic), representing the multidimensional convex hull volume occupied by the community, 2) Functional evenness (FEve) measuring the regularity, or evenness, of species in functional trait space weighted by their abundance, and 3) Functional divergence (FDiv), the distribution of abundance within the volume of functional trait space within each community (Villéger et al. 2008, Laliberté and Legendre 2010; see Appendix S1.4 for descriptions of the FD indices and their ecological meanings). FEve accounts for the relative abundance of species and is independent of species richness (Laliberté and Legendre 2010).

**Appendix S1.1.**
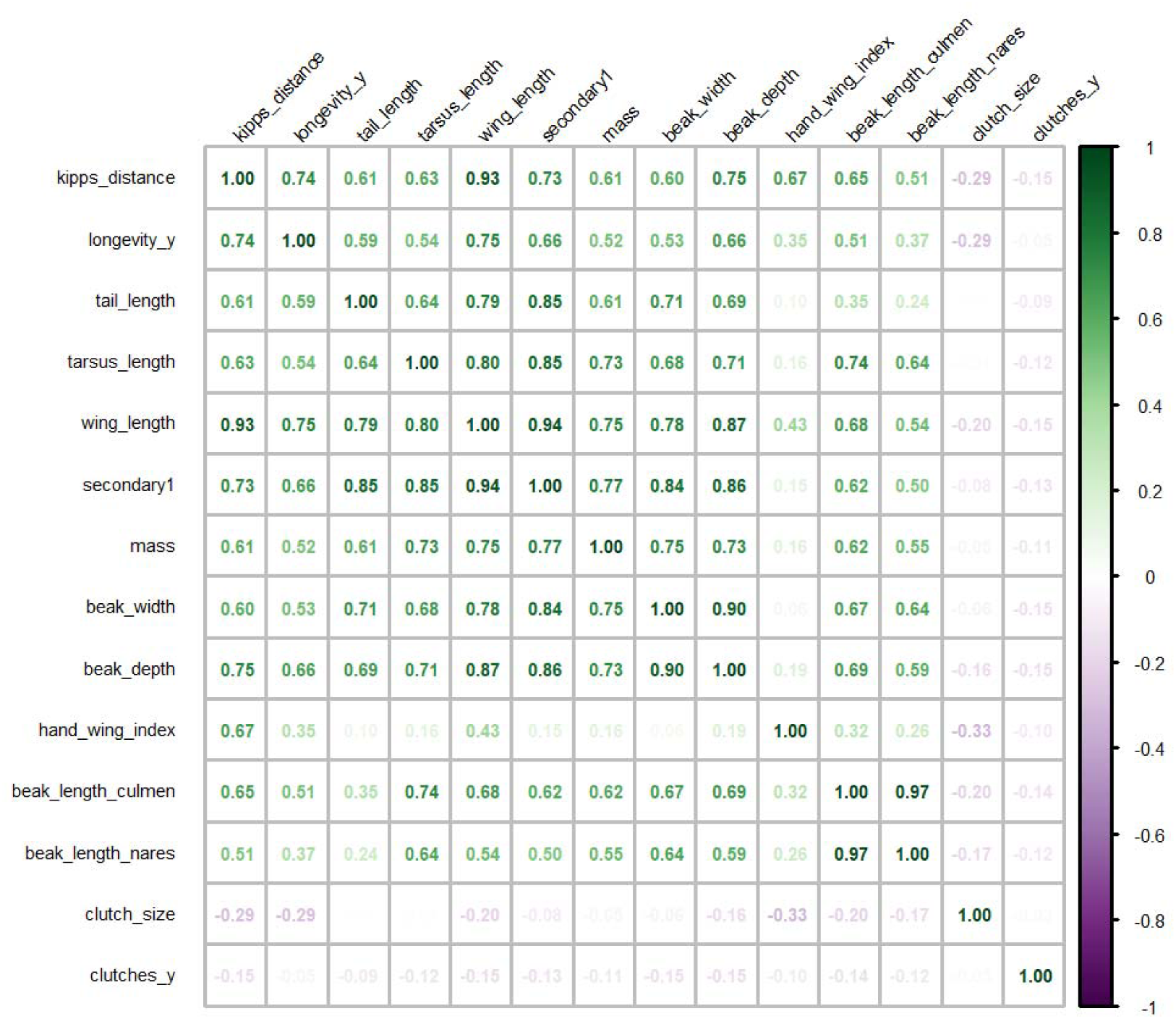
Correlation matrix of bird species traits considered for functional diversity calculations. The number depicts the Spearman correlation coefficient.

**Appendix S1.2.**
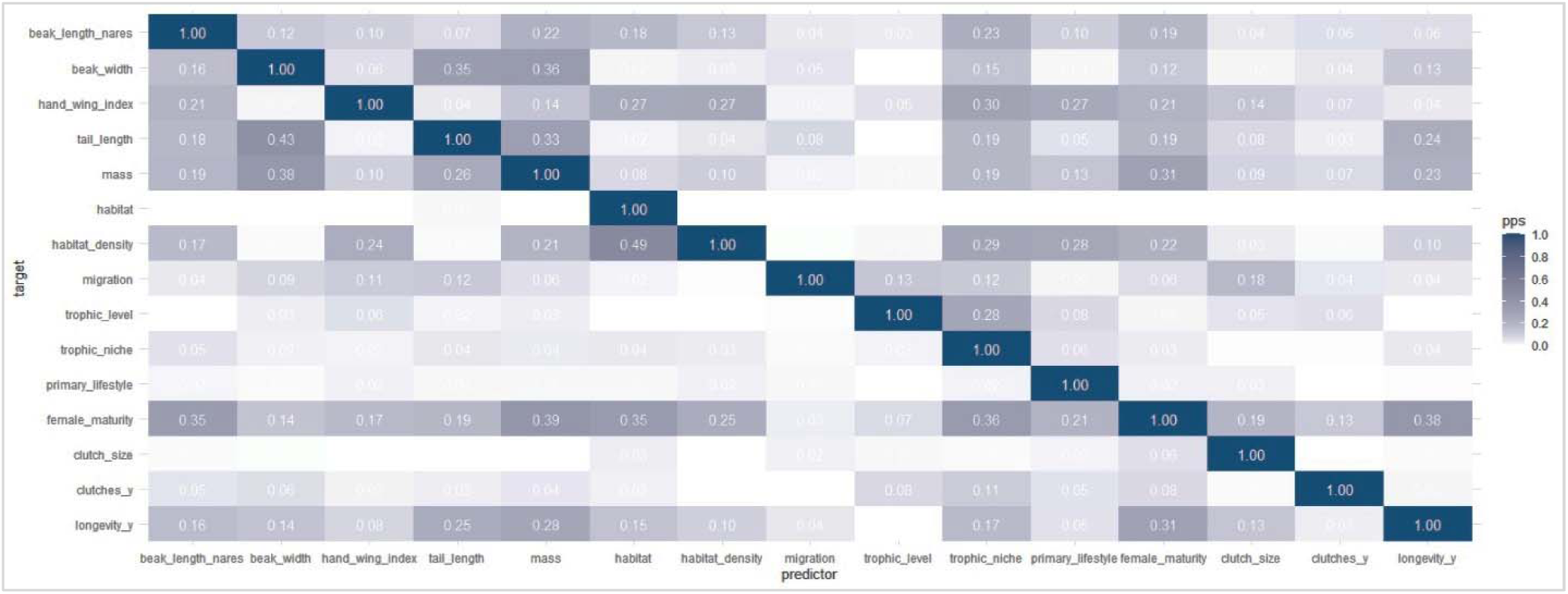
Predictive power score (PPS) matrix of the continuous and categorical bird species trait variables considered for functional diversity calculations. The number depicts the PPS, an asymmetric, data type-independent score that can detect linear or non-linear relationships between two columns with a value between 0 (no predictive power) and 1 (perfect predictive power).

**Appendix S1.3.** Bird species traits used to calculate functional diversity indices. The ecological functions of each trait and the variable type and source of the trait data are also described.

| Trait | Function | Variable type | Data source |
| --- | --- | --- | --- |
| Tarsus length | habitat use, foraging, movement, life history | continuous | AVONET<br>(Tobias et al., 2022) |
| Beak length nares | foraging, habitat use | continuous | AVONET |
| Beak depth | foraging, habitat use | continuous | AVONET |
| Hand-wing index | habitat use, movement | continuous | AVONET |
| Tail length | movement | continuous | AVONET |
| Habitat | habitat use | categorical | AVONET |
| Habitat density | habitat use | discrete | AVONET |
| Migration | movement, habitat use | discrete | AVONET |
| Trophic level | foraging, habitat use | categorical | AVONET |
| Trophic niche | foraging, habitat use | categorical | AVONET |
| Primary lifestyle | movement, habitat use | categorical | AVONET |
| Clutch size | life history | discrete | Amniote database<br>(Myhrvold et al., 2015) |
| Clutches per year | life history | discrete | Amniote database |
| Longevity | life history | discrete | Amniote database |

**Appendix S1.4.** Descriptions of functional diversity indices and their ecological meaning.

| Index | Description | Ecological meaning |
| --- | --- | --- |
| Functional richness<br>FRic | multidimensional convex hull volume occupied by the community (Laliberté & Legendre, 2010; Villéger et al., 2008) | The amount of functional niche space filled by species. An indicator of productivity, and resilience in the face of environmental fluctuations (Mason et al., 2005). |
| Functional evenness<br>FEve | regularity, or evenness, of species in functional trait space weighted by their abundance | High values represent regular/even trait distribution, and low values clumped/ uneven distributions in trait space. Lower values indicate that some parts of niche space are under-utilized. This can reduce productivity and reliability and increase opportunities for invaders (Mason et al., 2005). |
| Functional diversity<br>FDiv | distribution of abundance within the volume of functional trait space within each community | High values represent broad niches and indicate differences between species and thus low resource competition. Low values represent narrow niches and indicate similarities between species. Communities with low functional divergence may have lower ecosystem function because of less efficient resource use (Mason et al., 2005). |

**Appendix S1.5.**
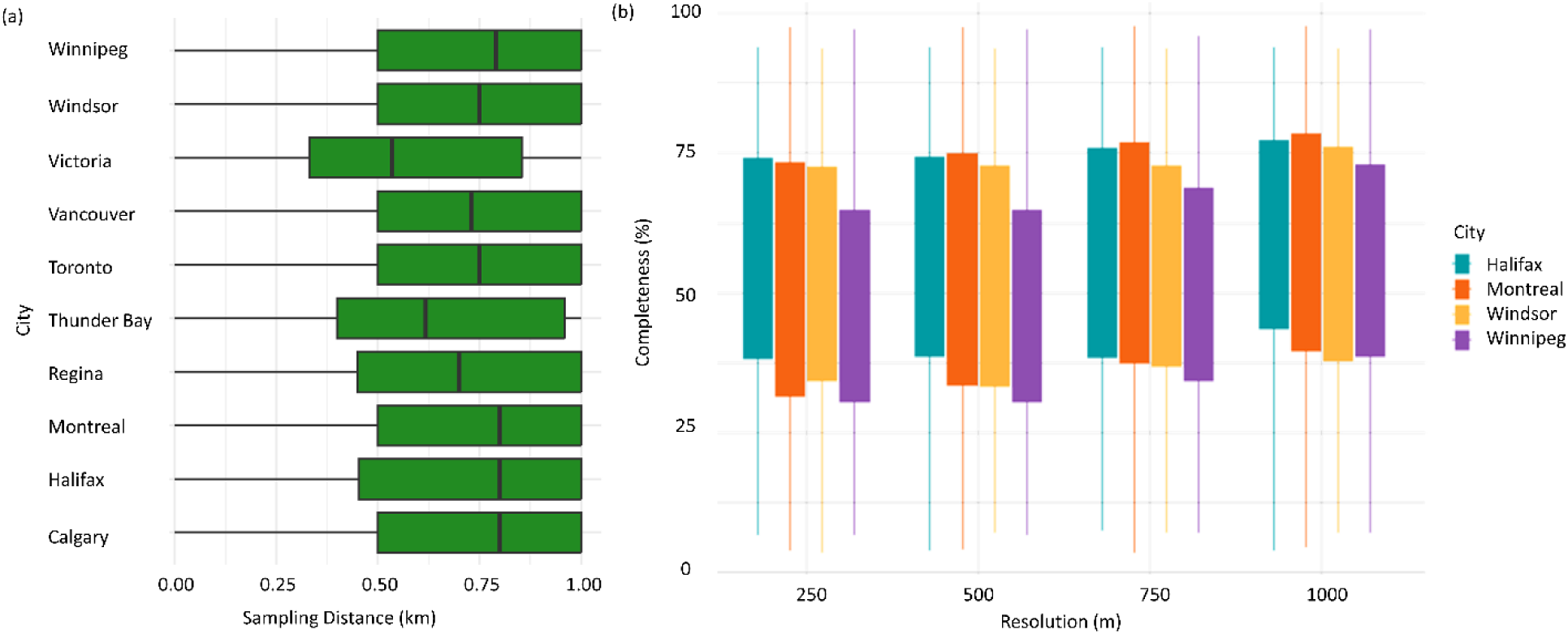
Barplots depicting a) variation in eBird bird sampling distance, measured as distance travelled during a birding occasion, across ten Canadian cities, and b) variation in sampling completeness (%) at four different grid resolutions (m) across four Canadian cities.

**Appendix S1.6.**
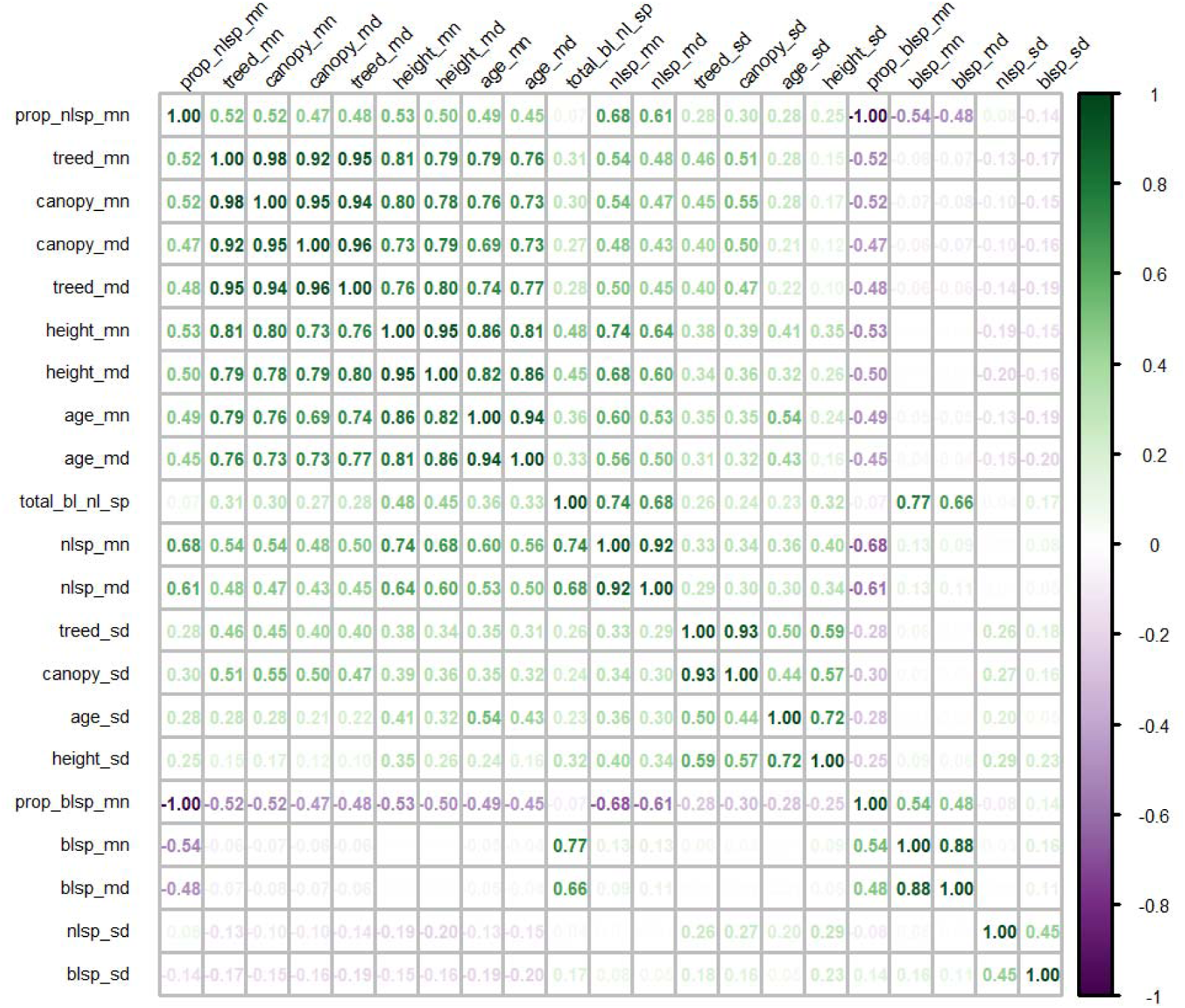
Correlation matrix of the vegetation traits considered as predictor variables for analysis. The number depicts the Spearman correlation coefficient.

**Appendix S1.7.**
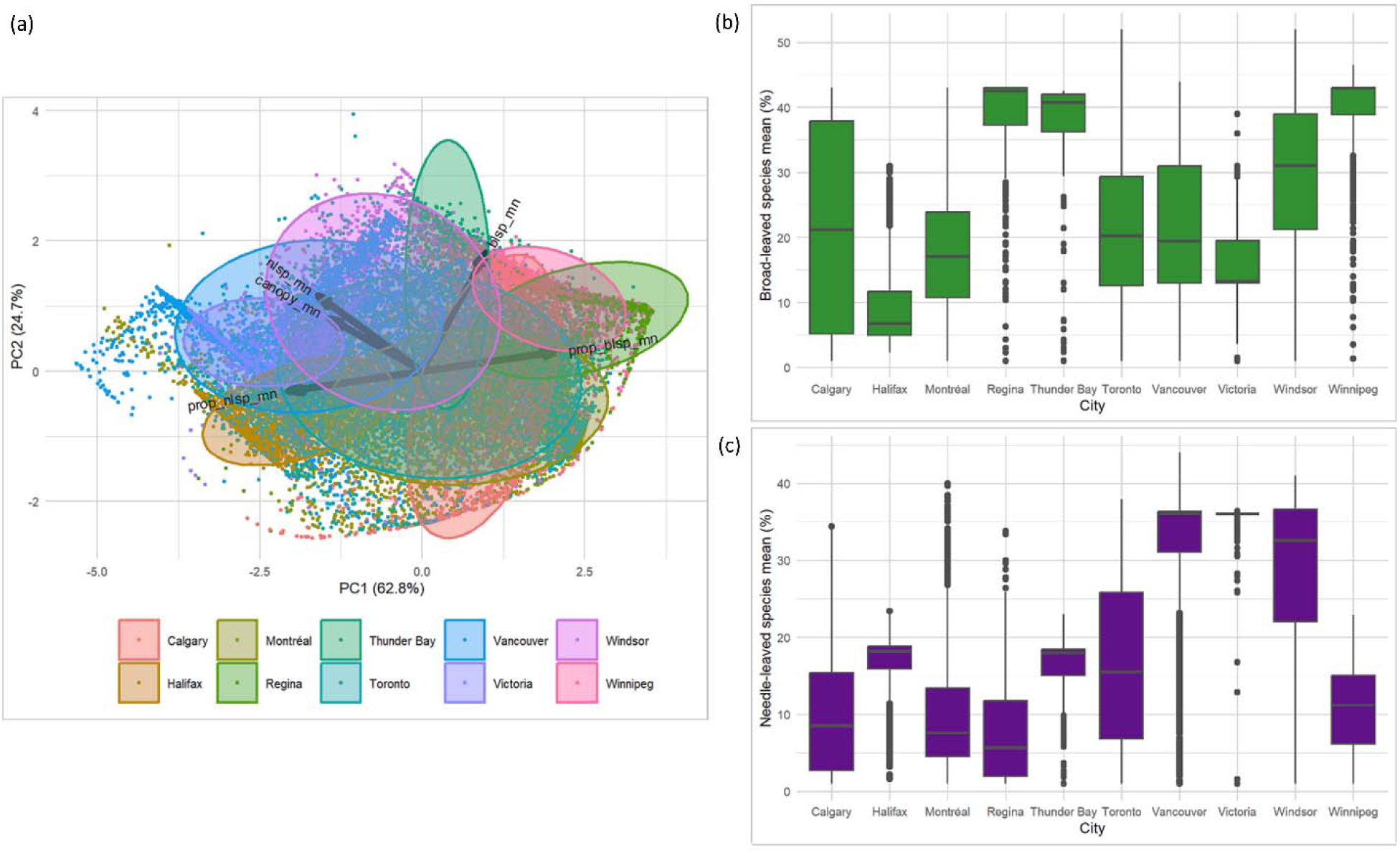
a) PCA-biplot showing the similarities and dissimilarities in vegetation between the ten Canadian cities. The plot shows the directionality of the vegetation attributes (black arrows) and the distribution of sampled points (colored according to city) on the first two principal components (PC1 and PC2). The colored data ellipses represent the differences in scores for each city. Barplots (on the right) depict the variation in the average b) broad-leaved species percentage and c) needle-leaved species percentage across the ten Canadian cities.

We re-sampled all vegetation variables to 750 m spatial resolution using the pixel aggregate method. We used R package *exactextractr* (version 0.9.1; Baston, 2020) and summarized carbon and vegetation raster values over city grids. For each polygon, we extracted the mean, sum and standard deviation of total forest carbon (kg C/m²) for each grid. We also extracted the mean, mode (the most common cell value) and standard deviation of each vegetation variable including canopy cover (percent crown closure), tree stand age (mean age of the leading species of polygons in years), stand height (mean height of the leading species in the polygons in meters), percentage of vegetated treed area (the proportion from vegetated area that is tree cover), and the percentage of coniferous and deciduous species (percent composition of all needle-leaf or broad-leaf species).

Vegetation structure variables suspected to be correlated underwent Spearman rank correlation analysis (Appendix S1.6). If two variables had a high (>70) correlation, we chose the most ecologically relevant variable for our analyses. We chose the mean and standard deviation (SD) of canopy cover, standard deviation of tree stand age, and standard deviation of tree height as our vegetation metrics. We also calculated the proportion of broad-leaved and needle-leaved species in each grid by first summing together the total percent composition of both (total trees = total broad-leaved + total needle-leaved species) and then dividing by the percent composition of each group.

### Appendix S2 Results

**Appendix S2.1.**
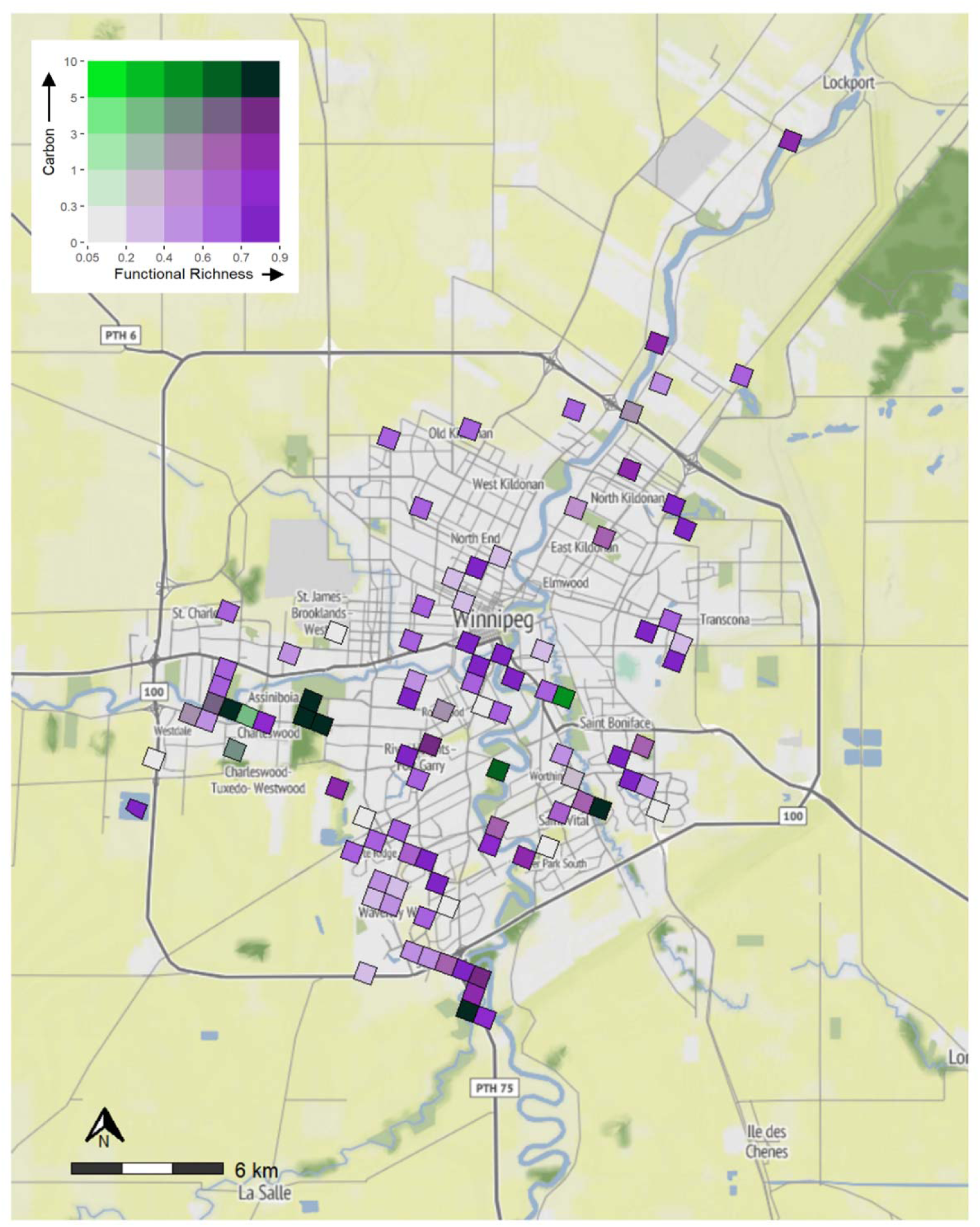
Bivariate map showing the spatial overlap between bird functional richness and above-ground forest carbon (kg C m^−2^) in 750 m grid cells across Winnipeg, Manitoba, Canada. Color scales display k-means clusters at 20% intervals. Dark green regions in the top right corner represent “hotspot” areas with the highest overlapping (the top 20%) values for functional richness and total carbon. Light grey areas in the bottom left corner represent cold spots, with the lowest (bottom 20%) values for both variables. Dark purple regions in the bottom right and bright green regions in the top left represent trade-off areas or grids with the highest functional richness and low carbon values or vice versa.

**Appendix S2.2.**
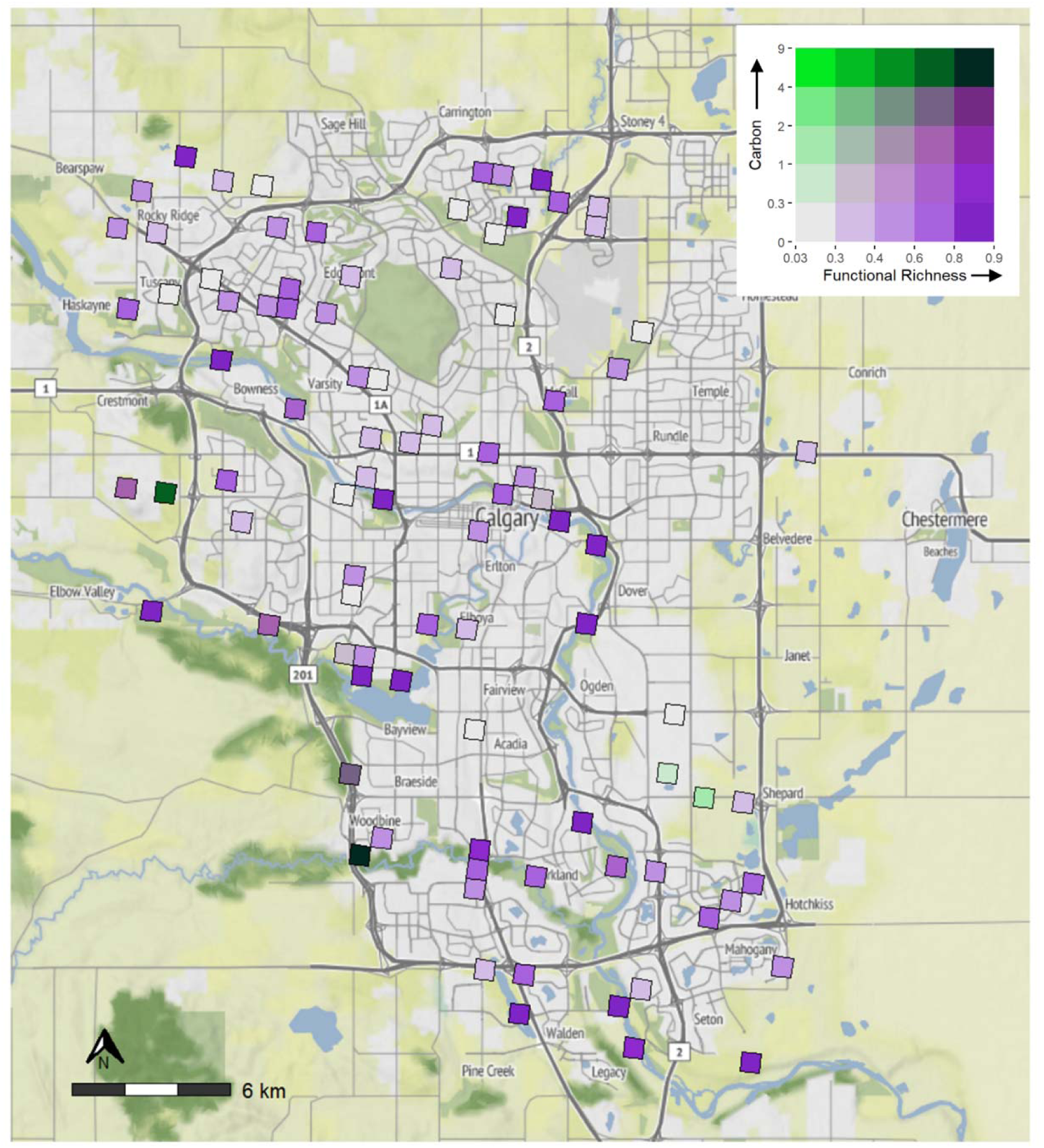
Bivariate map showing the spatial overlap between bird functional richness and above-ground forest carbon (kg C m^−2^) in 750 m grid cells across Calgary, Alberta, Canada. Color scales display k-means clusters at 20% intervals. Dark green regions in the top right corner represent “hotspot” areas with the highest overlapping (the top 20%) values for functional richness and total carbon. Light grey areas in the bottom left corner represent cold spots, with the lowest (bottom 20%) values for both variables. Dark purple regions in the bottom right and bright green regions in the top left represent trade-off areas or grids with the highest functional richness and low carbon values or vice versa.

**Appendix S2.3.**
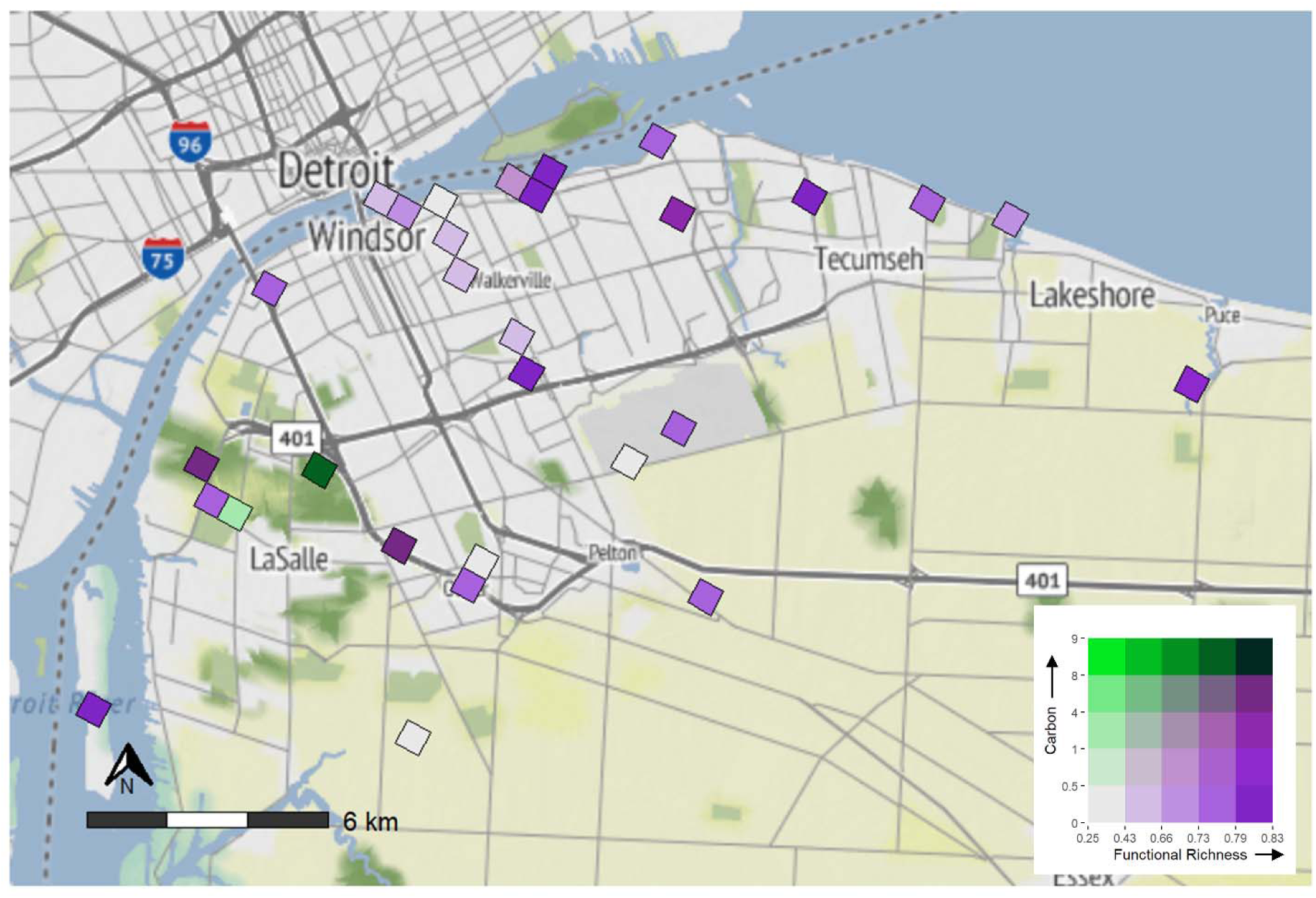
Bivariate map showing the spatial overlap between bird functional richness and above-ground forest carbon (kg C m^−2^) in 750 m grid cells across Windsor, Ontario, Canada. Color scales display k-means clusters at 20% intervals. Dark green regions in the top right corner represent “hotspot” areas with the highest overlapping (the top 20%) values for functional richness and total carbon. Light grey areas in the bottom left corner represent cold spots, with the lowest (bottom 20%) values for both variables. Dark purple regions in the bottom right and bright green regions in the top left represent trade-off areas or grids with the highest functional richness and low carbon values or vice versa.

**Appendix S2.4.**
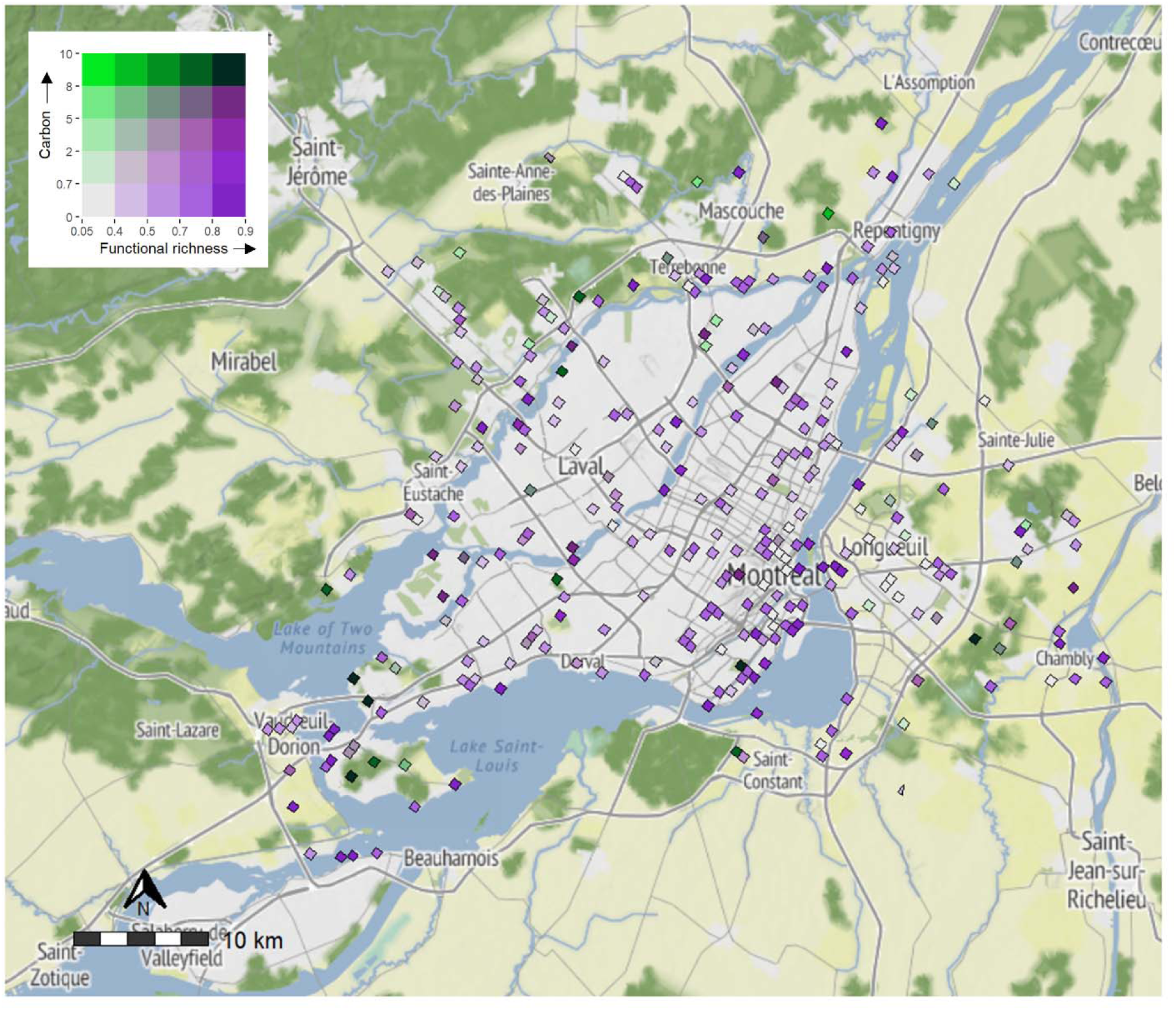
Bivariate map showing the spatial overlap between bird functional richness and above-ground forest carbon (kg C m^−2^) in 750 m grid cells across Montreal, Quebec, Canada. Color scales display k-means clusters at 20% intervals. Dark green regions in the top right corner represent “hotspot” areas with the highest overlapping (the top 20%) values for functional richness and total carbon. Light grey areas in the bottom left corner represent cold spots, with the lowest (bottom 20%) values for both variables. Dark purple regions in the bottom right and bright green regions in the top left represent trade-off areas or grids with the highest functional richness and low carbon values or vice versa.

**Appendix S2.5.**
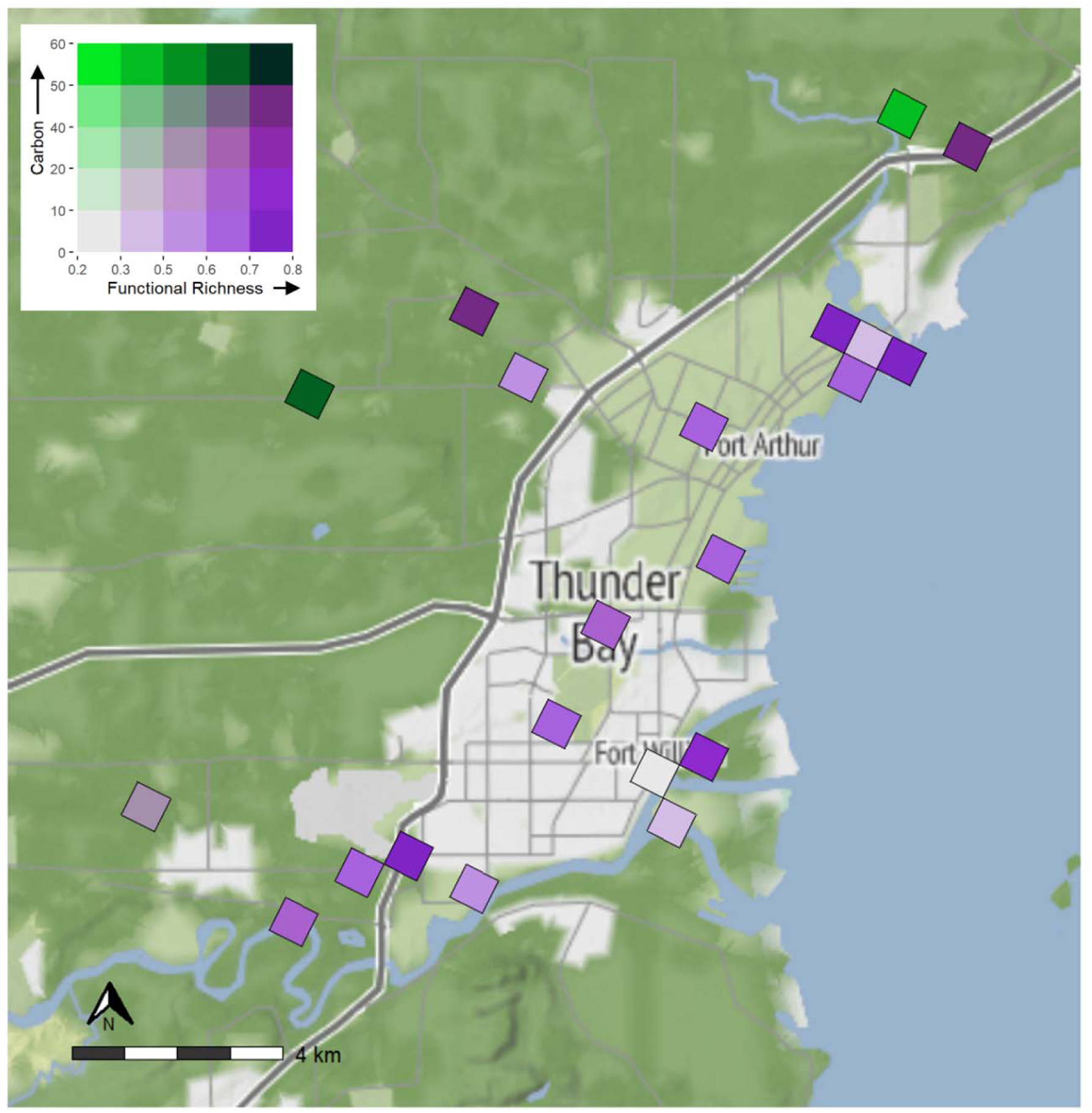
Bivariate map showing the spatial overlap between bird functional richness and above-ground forest carbon (kg C m^−2^) in 750 m grid cells across Thunder Bay, Ontario, Canada. Color scales display k-means clusters at 20% intervals. Dark green regions in the top right corner represent “hotspot” areas with the highest overlapping (the top 20%) values for functional richness and total carbon. Light grey areas in the bottom left corner represent cold spots, with the lowest (bottom 20%) values for both variables. Dark purple regions in the bottom right and bright green regions in the top left represent trade-off areas or grids with the highest functional richness and low carbon values or vice versa.

**Appendix S2.6.**
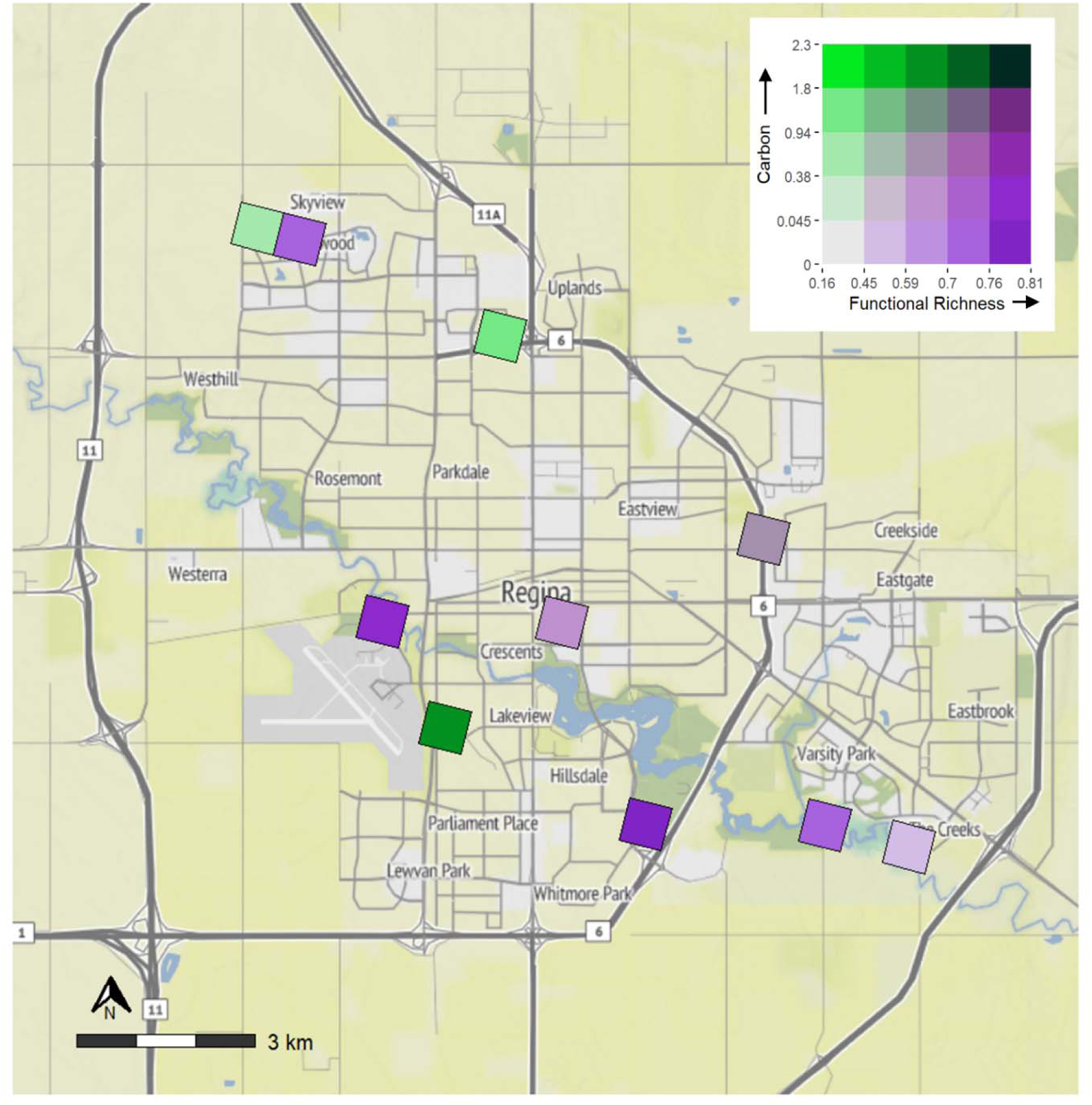
Bivariate map showing the spatial overlap between bird functional richness and above-ground forest carbon (kg C m^−2^) in 750 m grid cells across Regina, Saskatchewan, Canada. Color scales display k-means clusters at 20% intervals. Dark green regions in the top right corner represent “hotspot” areas with the highest overlapping (the top 20%) values for functional richness and total carbon. Light grey areas in the bottom left corner represent cold spots, with the lowest (bottom 20%) values for both variables. Dark purple regions in the bottom right and bright green regions in the top left represent trade-off areas or grids with the highest functional richness and low carbon values or vice versa.

**Appendix S2.7.**
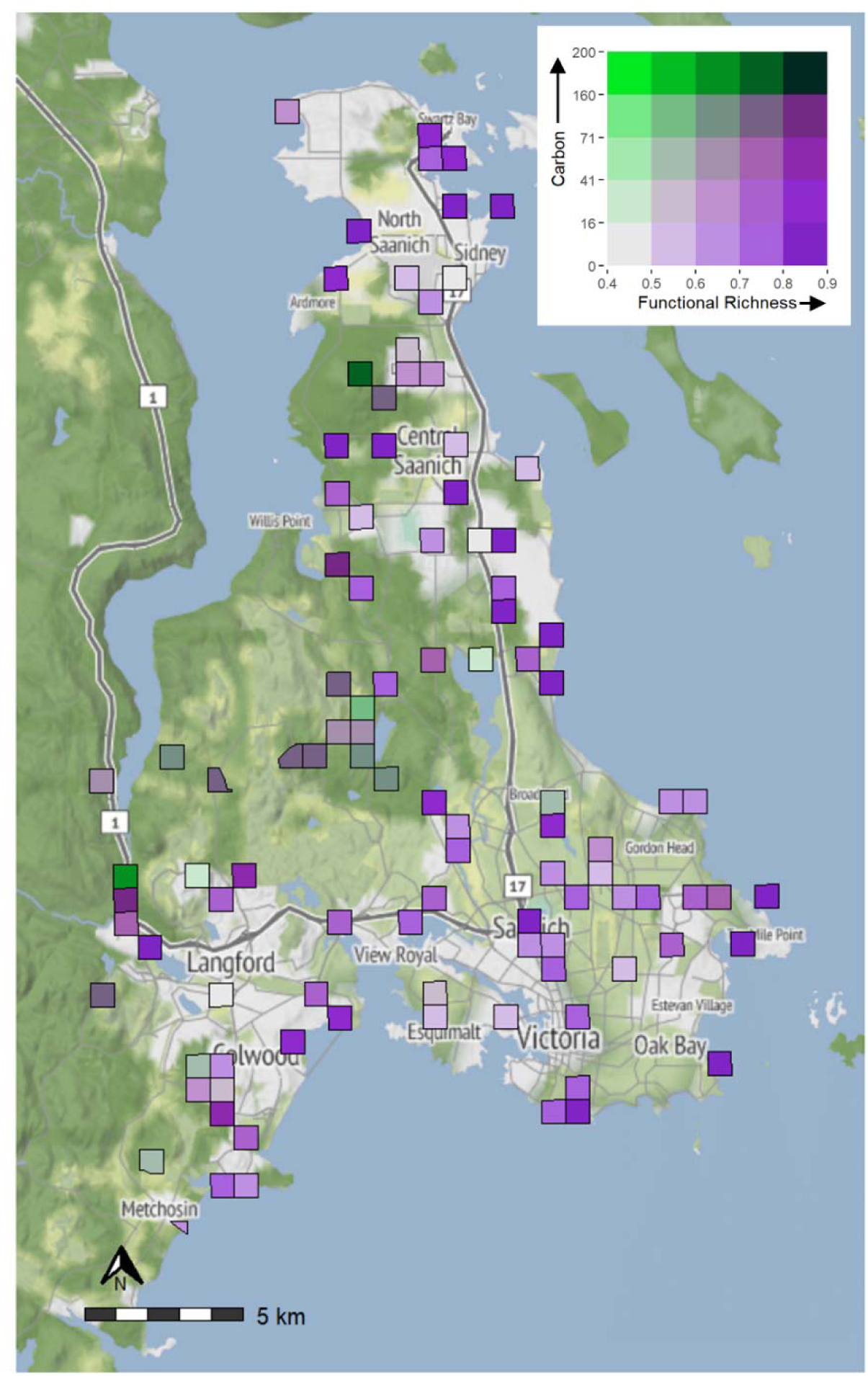
Bivariate map showing the spatial overlap between bird functional richness and above-ground forest carbon (kg C m^−2^) in 750 m grid cells across Victoria, British Columbia, Canada. Color scales display k-means clusters at 20% intervals. Dark green regions in the top right corner represent “hotspot” areas with the highest overlapping (the top 20%) values for functional richness and total carbon. Light grey areas in the bottom left corner represent cold spots, with the lowest (bottom 20%) values for both variables. Dark purple regions in the bottom right and bright green regions in the top left represent trade-off areas or grids with the highest functional richness and low carbon values or vice versa.

**Appendix S2.8.**
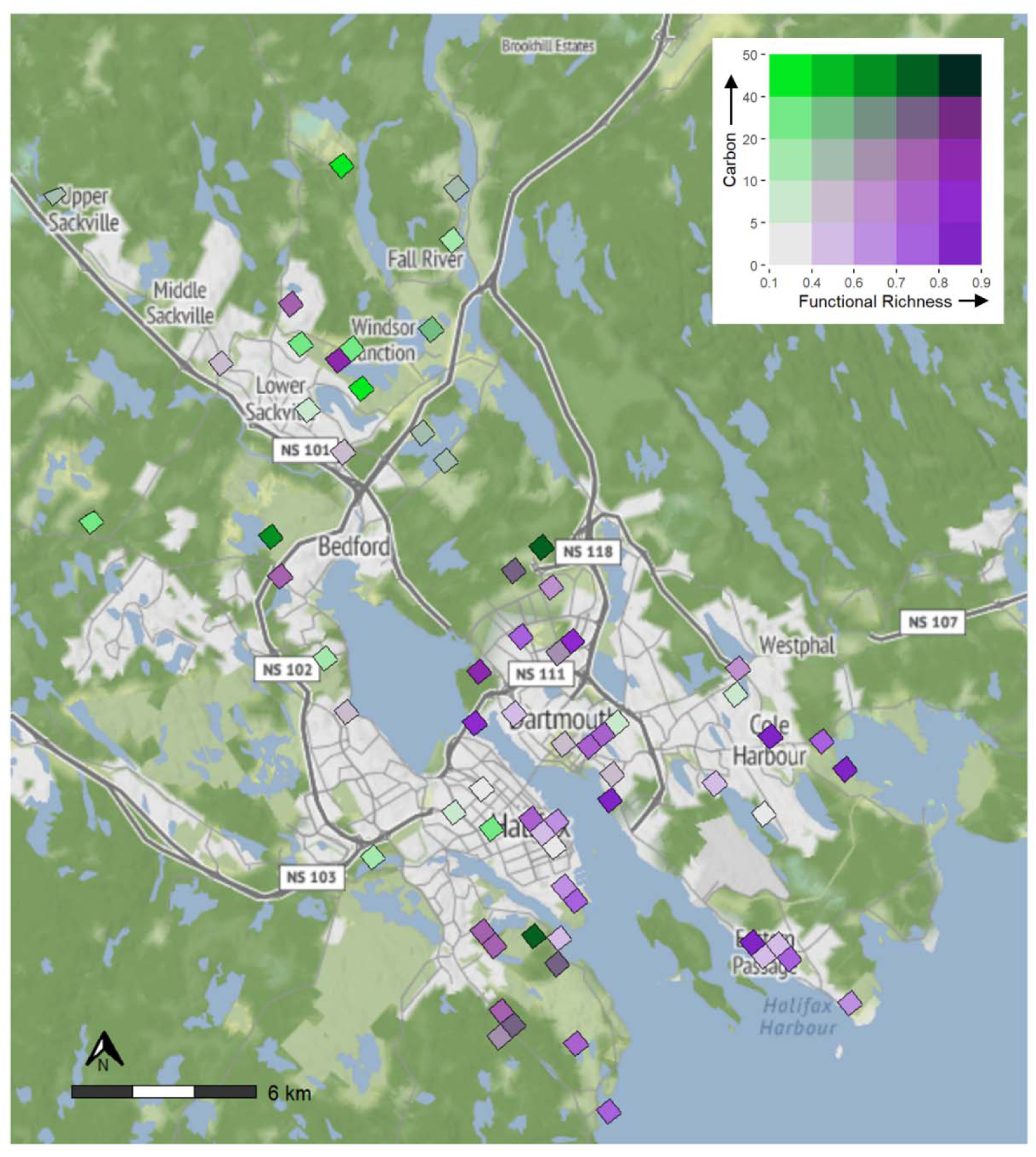
Bivariate map showing the spatial overlap between bird functional richness and above-ground forest carbon (kg C m^−2^) in 750 m grid cells across Halifax, Nova Scotia, Canada. Color scales display k-means clusters at 20% intervals. Dark green regions in the top right corner represent “hotspot” areas with the highest overlapping (the top 20%) values for functional richness and total carbon. Light grey areas in the bottom left corner represent cold spots, with the lowest (bottom 20%) values for both variables. Dark purple regions in the bottom right and bright green regions in the top left represent trade-off areas or grids with the highest functional richness and low carbon values or vice versa.

**Appendix S2.9.**
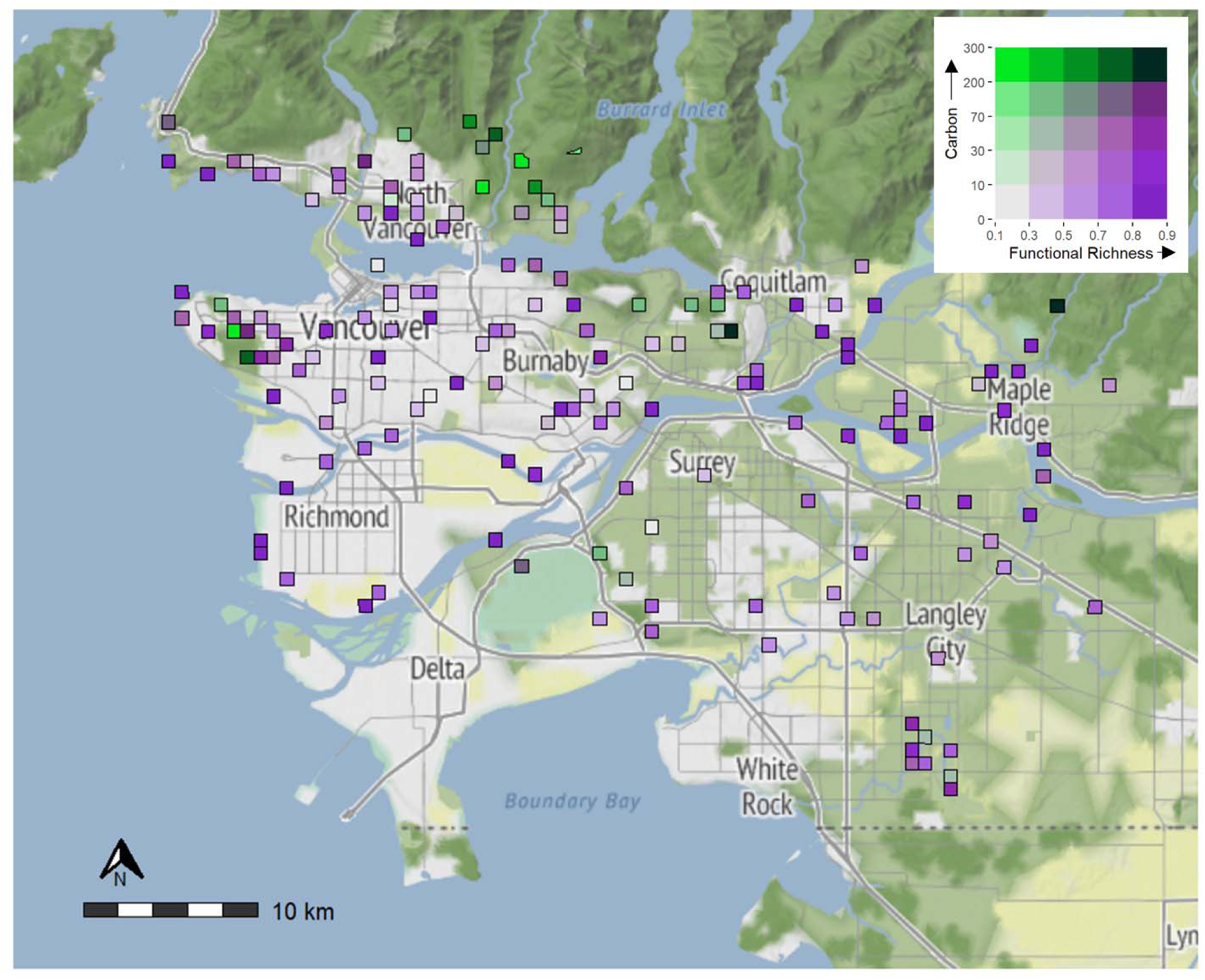
Bivariate map showing the spatial overlap between bird functional richness and above-ground forest carbon (kg C m^−2^) in 750 m grid cells across Vancouver, British Columbia, Canada. Color scales display k-means clusters at 20% intervals. Dark green regions in the top right corner represent “hotspot” areas with the highest overlapping (the top 20%) values for functional richness and total carbon. Light grey areas in the bottom left corner represent cold spots, with the lowest (bottom 20%) values for both variables. Dark purple regions in the bottom right and bright green regions in the top left represent trade-off areas or grids with the highest functional richness and low carbon values or vice versa.

**Appendix S2.10.**
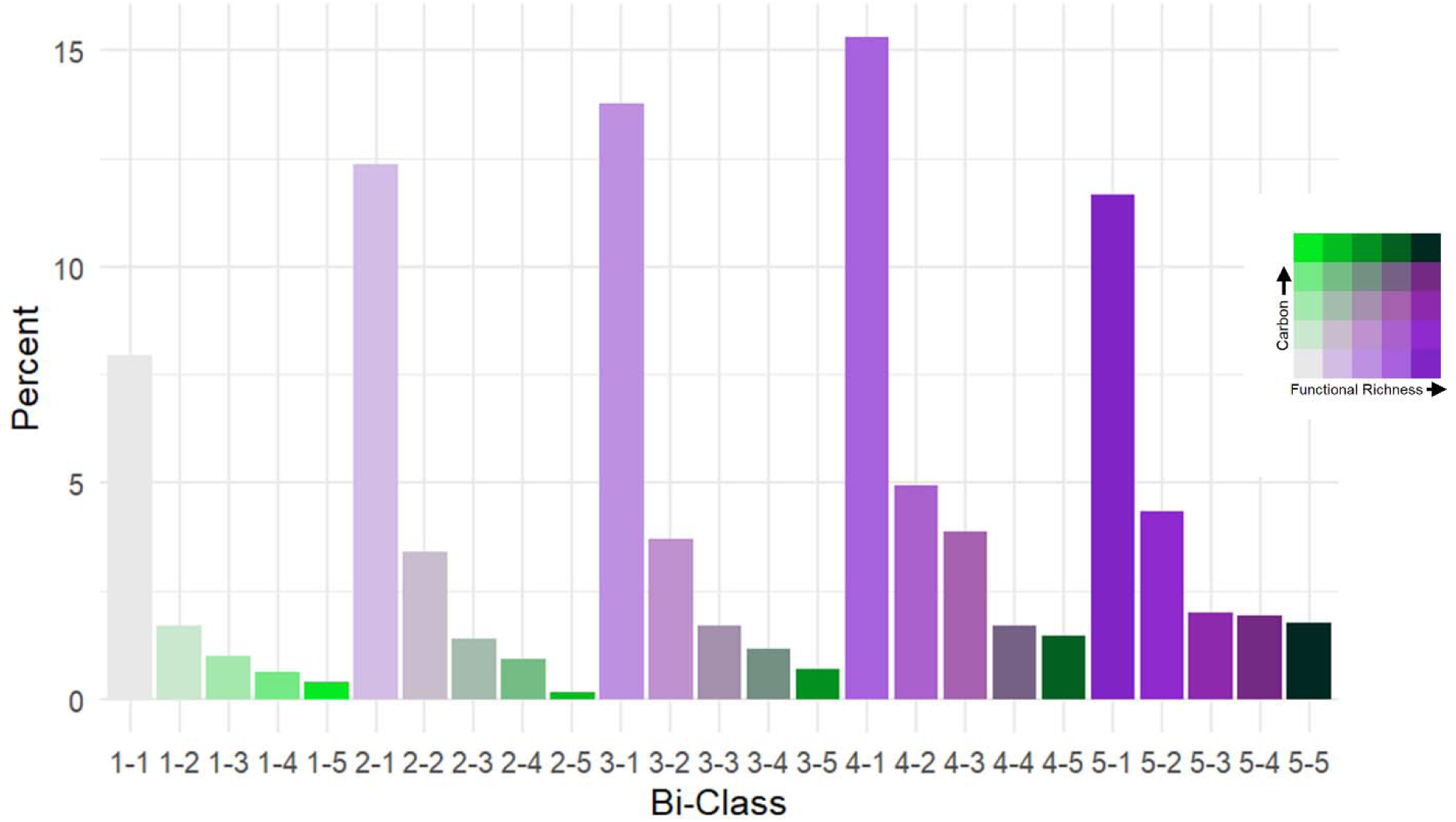
Barplot showing bivariate bi-class percentages between carbon and bird functional richness for ten Canadian cities. We calculated k-means clusters at 20% intervals for the functional richness and total carbon map layers for each city and re-classified cell values for each layer with a value of 1–5 based on which percentile range they fell into. The cell values were calculated using unique combinations of the two layers, which resulted in 25 different cell values (or bi-classes) used to produce bivariate maps for bird biodiversity by total forest carbon.

**Appendix S2.11.**
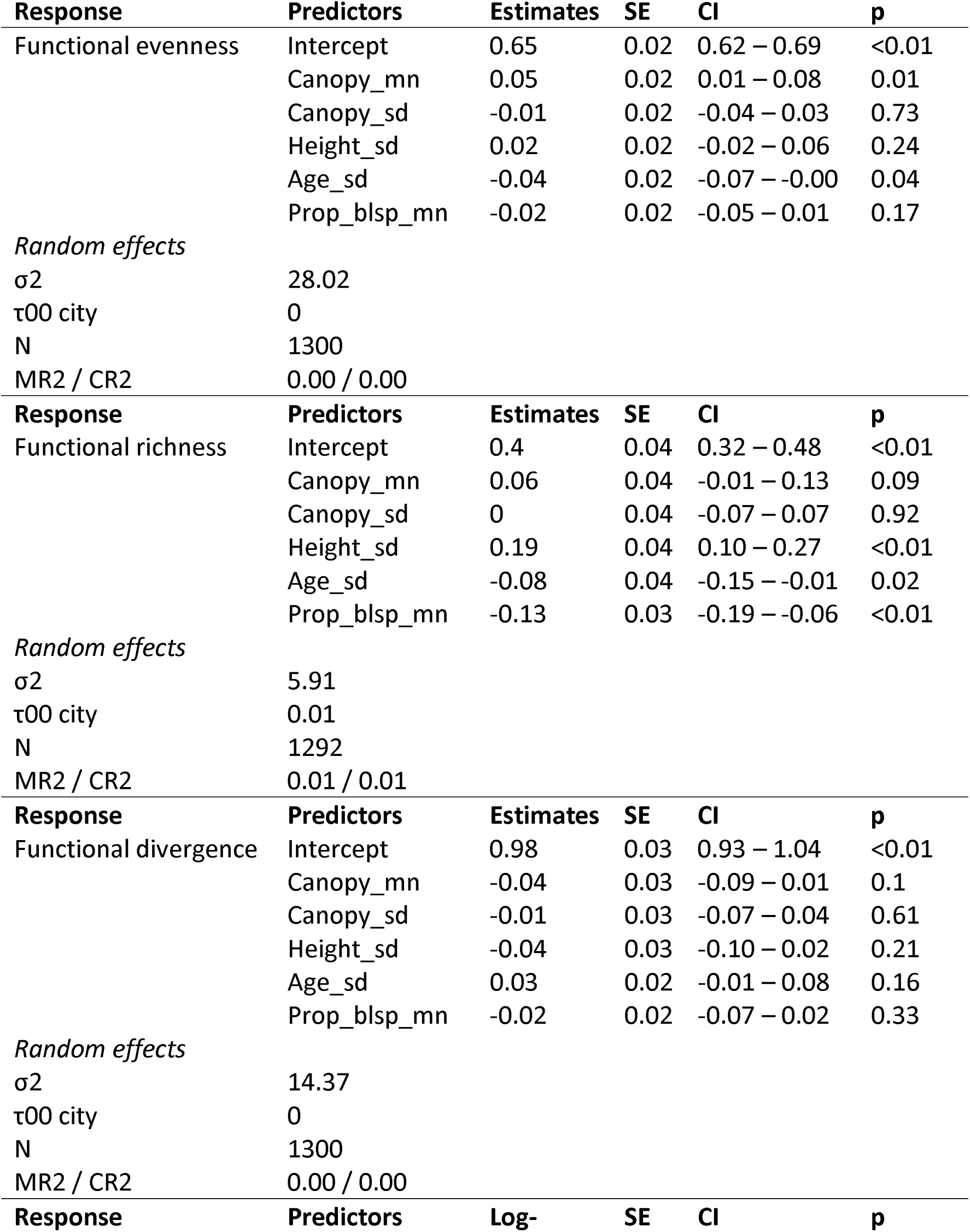
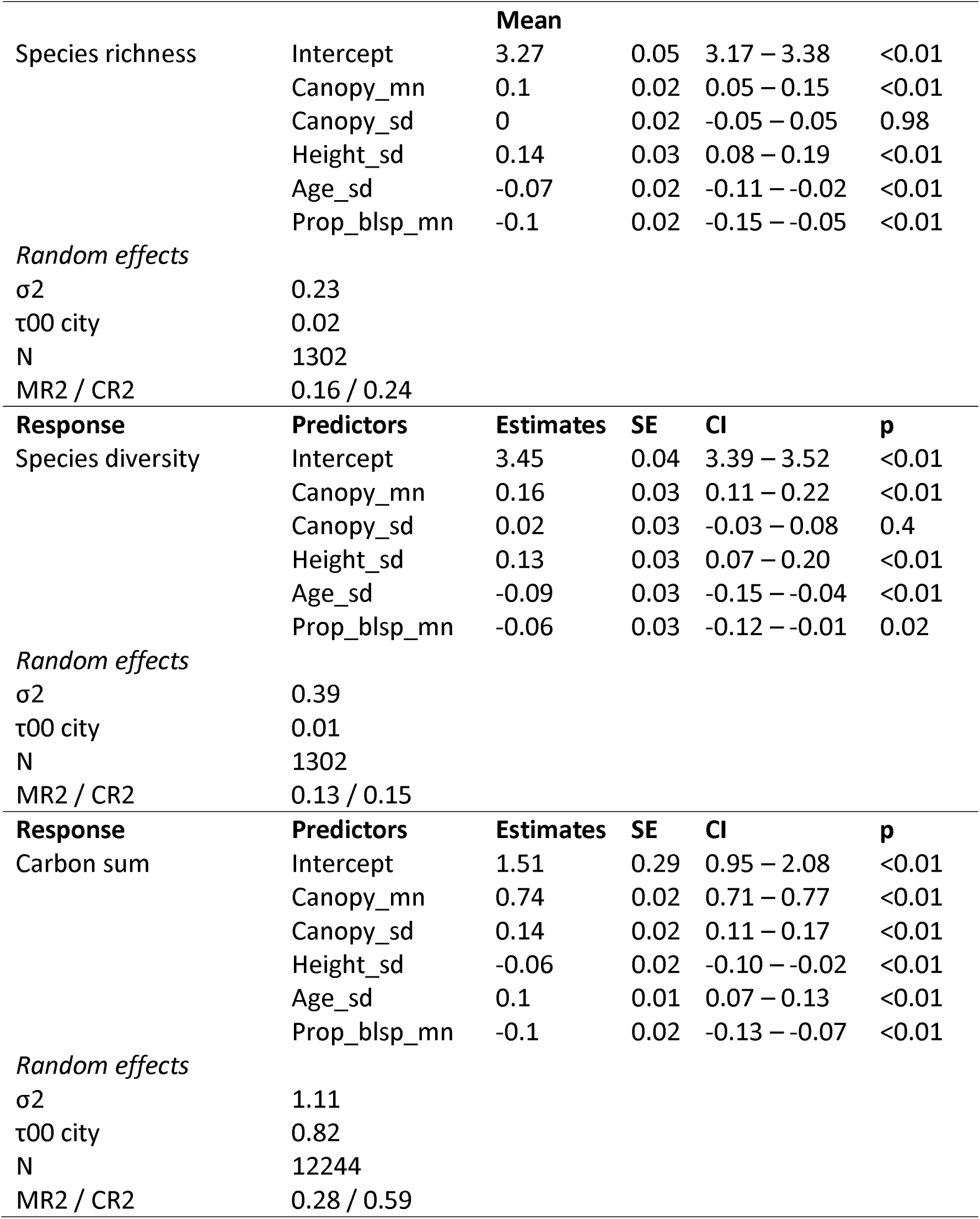
Generalized linear mixed model (GLMM) estimates, standard errors (SE), 95% confidence intervals and p-values for all cities combined, with city as a random effect. The symbol σ2 is the residual variance; τ00 is the variance among the random effects; N is the total number of observations; and MR2 / CR2 are the marginal and conditional R-squared values.

**Appendix S2.12.**
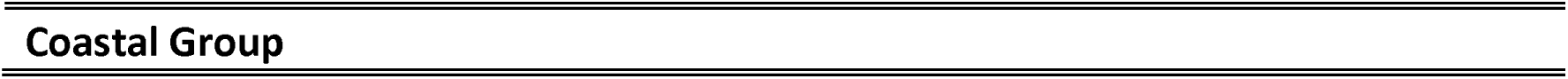
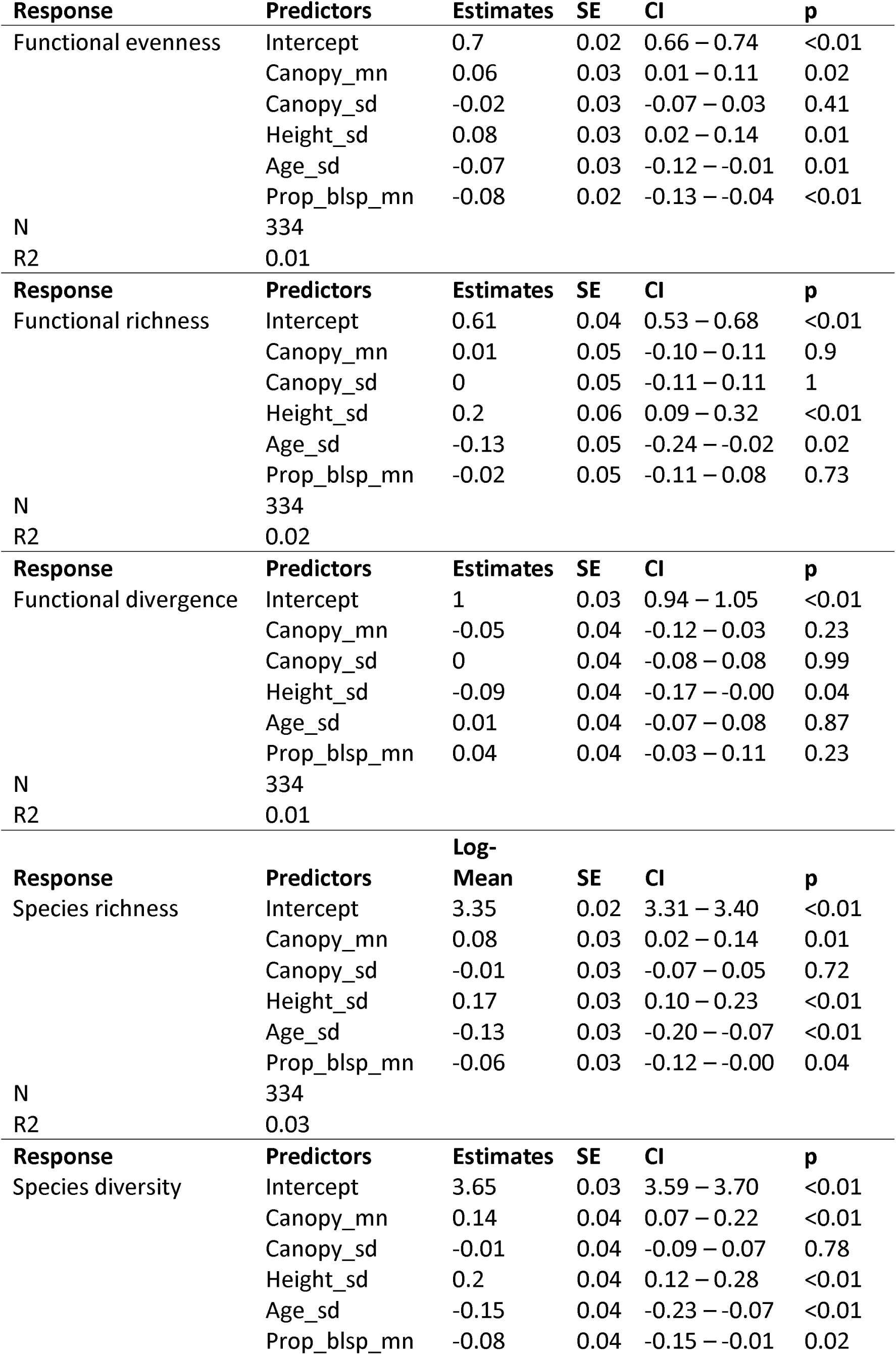
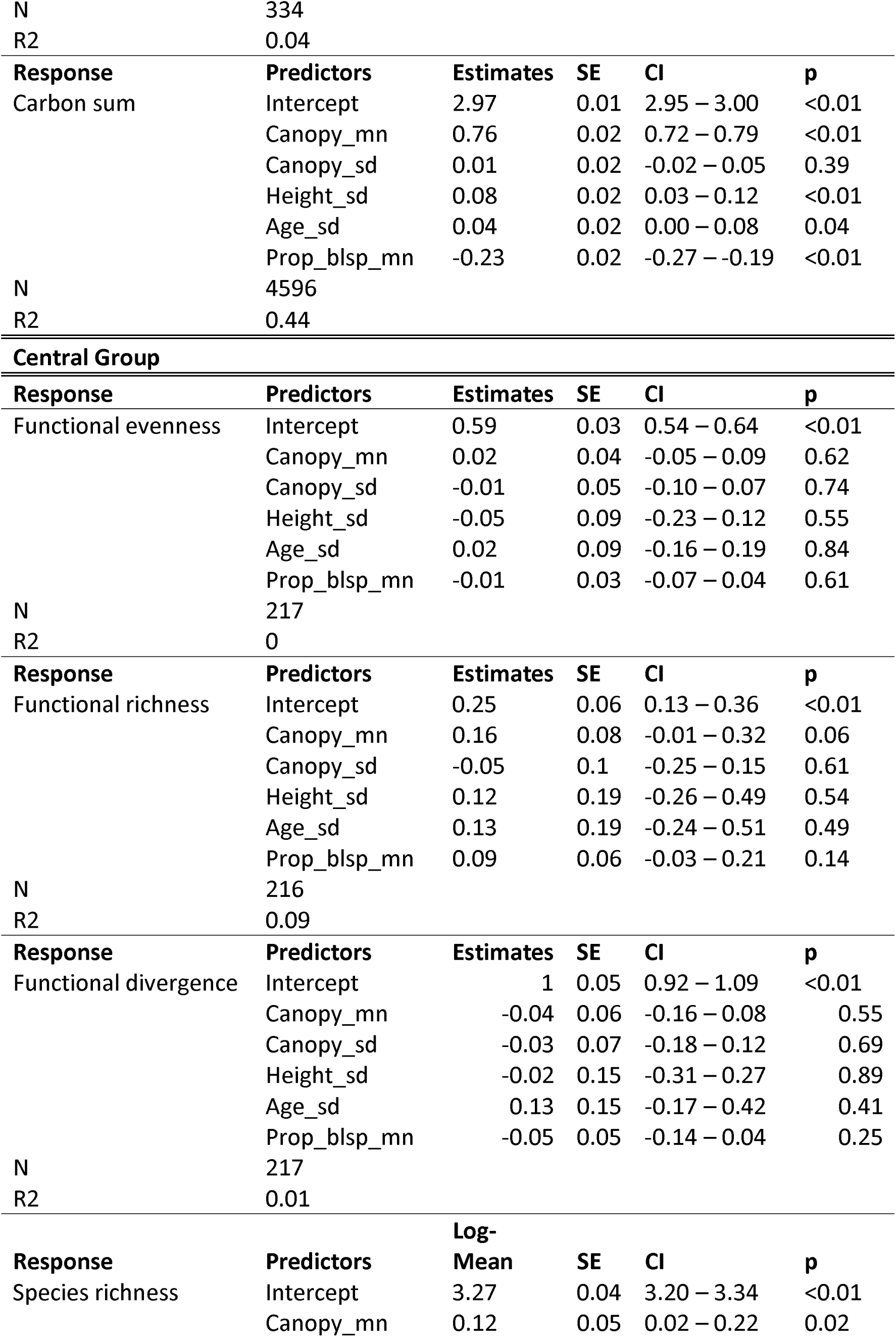
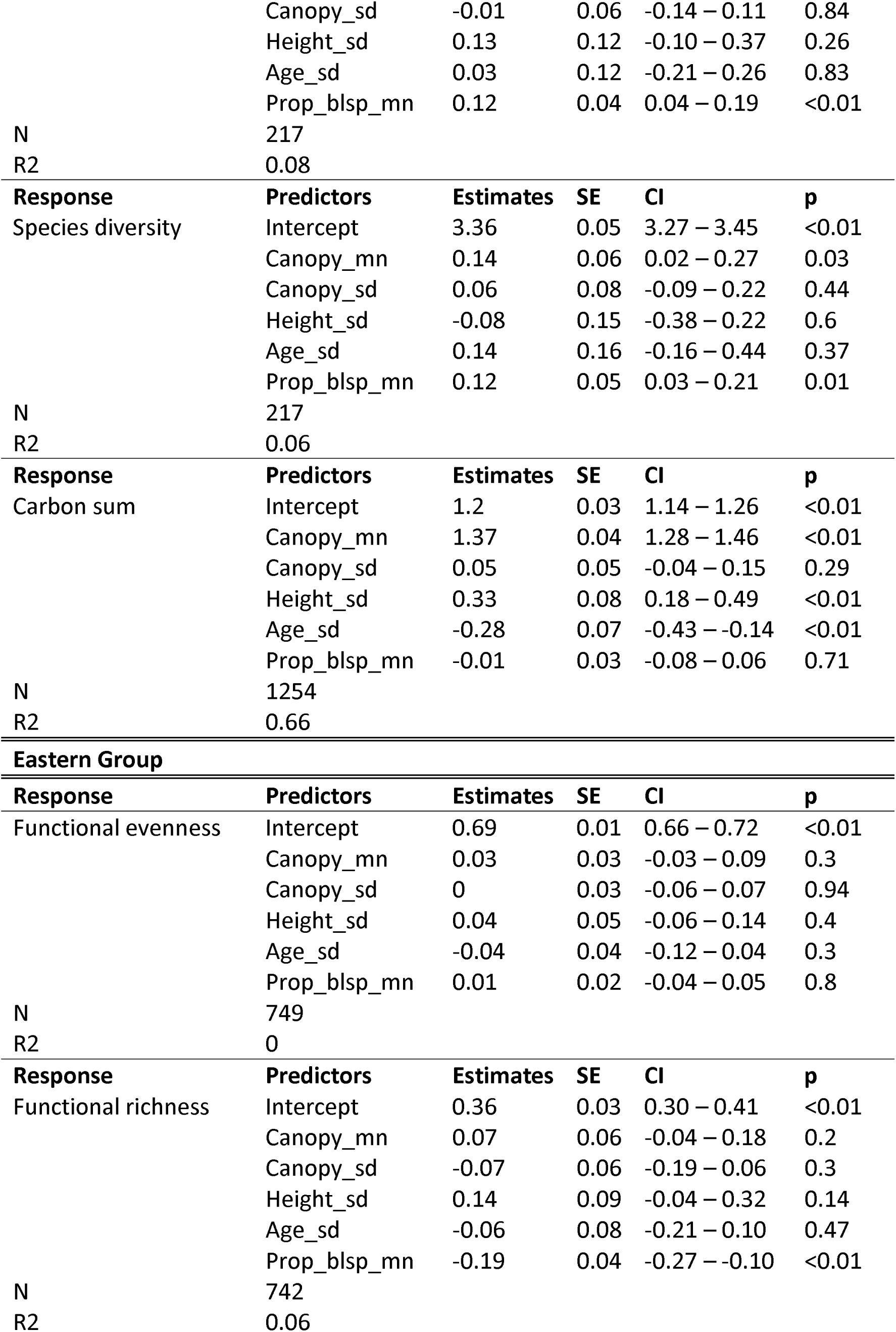
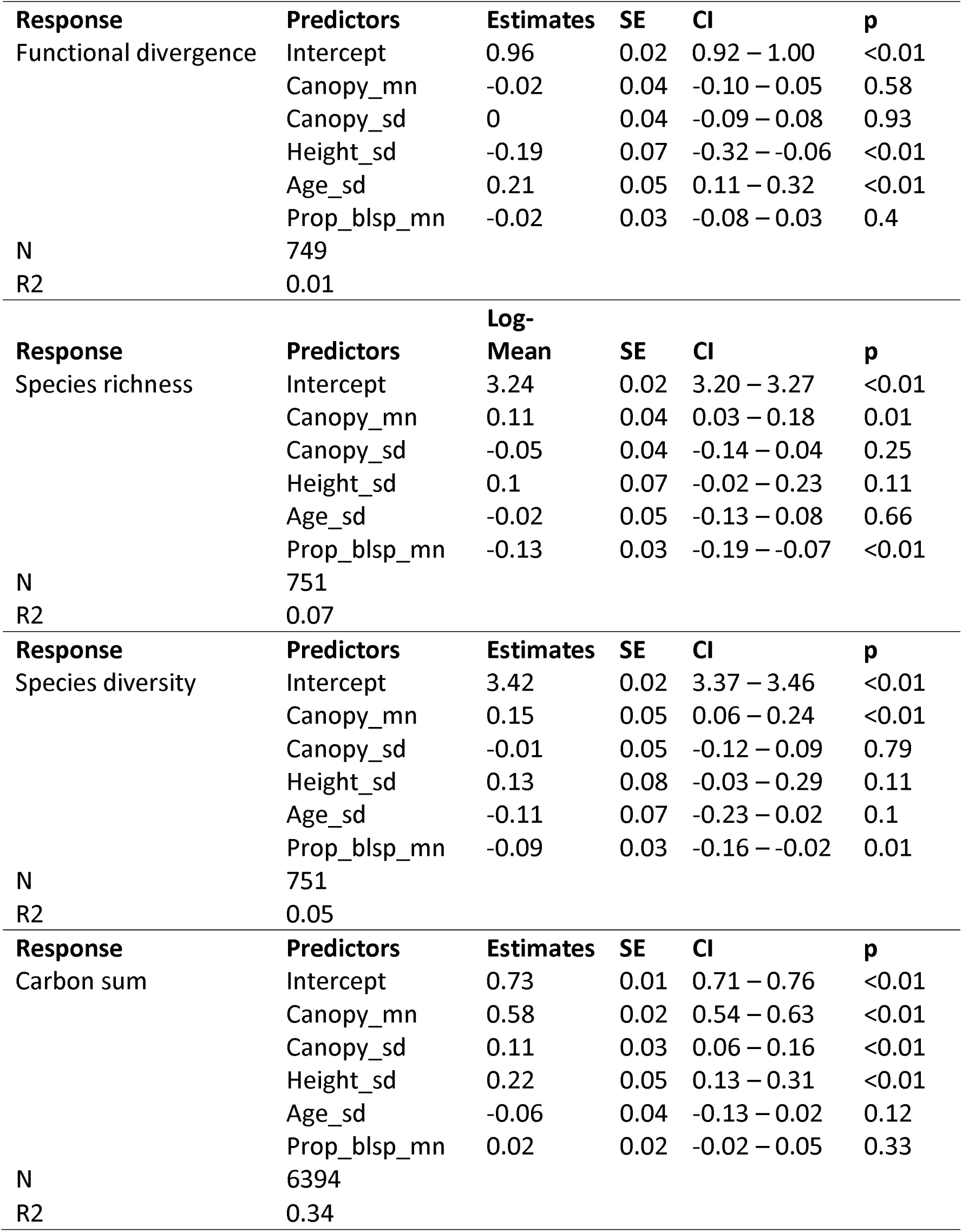
Generalized linear model estimates, standard errors (SE), 95% confidence intervals (CI) and p-values for the three different city groups. N is the total number of observations, and R2 is pseudo-R-squared.

**Figure S2.13.**
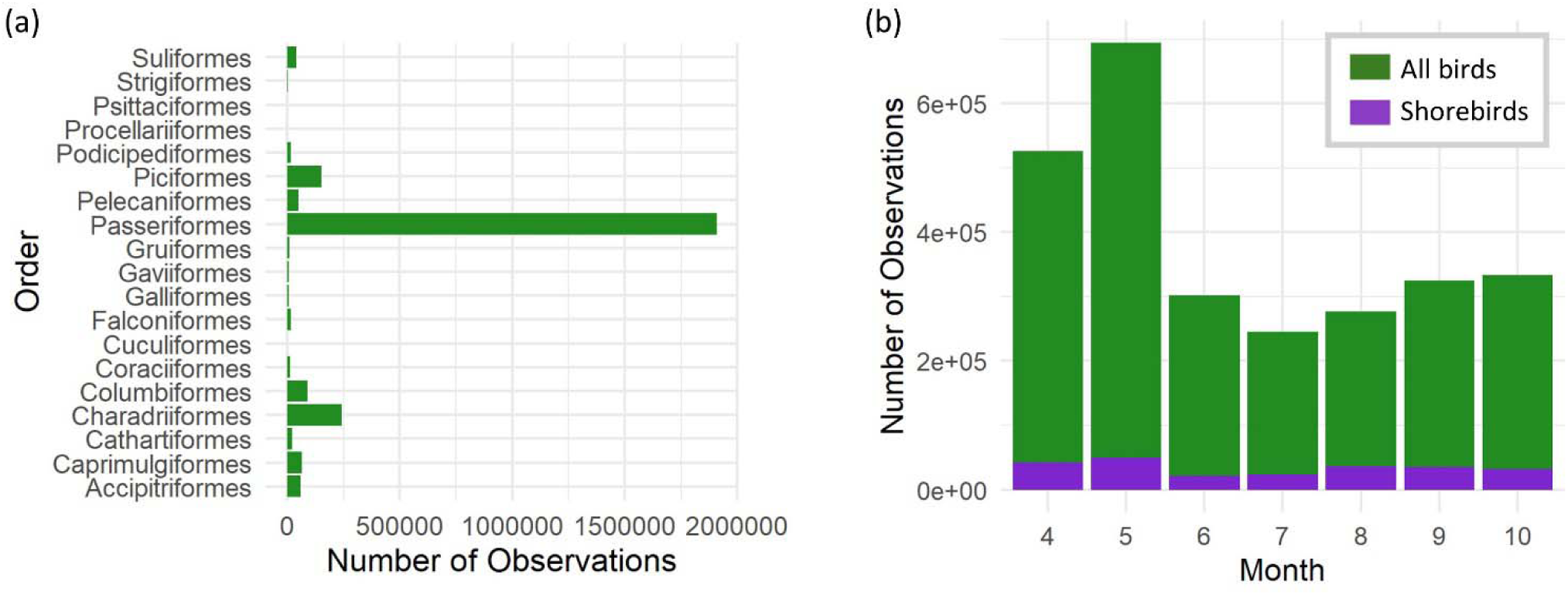
Barplots showing a) the number of total bird observations per taxonomic order, and b) the number of observations per month (during the years 2011-2021) for all bird orders, and shorebirds (Charadriiformes).

